# Assessing the oviposition and larval ecologies of mosquitoes and *Stegomyia* indices along an urban-rural gradient during ongoing dengue outbreaks in the arboviral hotspots of Cocody-Bingerville, southeastern Côte d’Ivoire

**DOI:** 10.1101/2025.01.02.631058

**Authors:** Yasmine N. Biré, Julien Z. B. Zahouli, Jean D. K. Dibo, Pierre N. Coulibaly, Prince G. Manouana, Jacques F. Mavoungou, Fanny Hellhammer, Gäel D. Maganga, Luc S. Djogbenou, Ayola A. Adegnika, Steffen Borrmann, Stefanie C. Becker, Mahama Touré

**Affiliations:** Centre d’Entomologie Médicale et Vétérinaire, Université Alassane Ouattara, Bouaké, Côte d’Ivoire; Centre Suisse de Recherches Scientifiques en Côte d’Ivoire, Abidjan, Côte d’Ivoire; Centre de Recherches Médicales de Lambaréné, Lambaréné, Gabon; Institut for Tropical Medicine, University of Tübingen, Tübingen, Germany; Institut de Recherche en Ecologie Tropicale, Centre National de la Recherche Scientifique et Technologique, Libreville, Gabon; Institute for Parasitology University of Veterinary Medicine Hanover, Hanover, Germany; Research Center for Emerging Infection and Zoonoses, University of Veterinary Medicine Hanover, Hanover, Germany; Centre Interdisciplinaire de Recherches Médicales de Franceville, Franceville, Gabon; Tropical Infectious Diseases Research Centre, University of Abomey-Calavi, Cotonou, Benin; German Center for Infection Research, Tübingen, Germany

**Keywords:** Arbovirus, *Aedes*, *Aedes aegypti*, Egg-laying, Breeding site, Predatory larvae, Ovitrap index, Larval index, Arboviral risk, Ecozone, Urbanization, Africa

## Abstract

**Background:** Côte d’Ivoire (West Africa) has recurrently faced multiple outbreaks of *Aedes* mosquito-transmitted arboviral diseases (e.g., dengue (DEN) and yellow fever (YF)), mainly in the health district of Cocody-Bingerville subjected to rapid urbanization. Thus, this study assessed the ecology of mosquito immatures and *Stegomyia* indices along an urban-rural gradient during ongoing dengue outbreaks within arboviral hotspots in Cocody-Bingerville.

**Methods:** Between August 2023 and July 2024, we collected mosquito eggs and larvae in urban, suburban and rural areas of Cocody-Bingerville using standard ovitraps and larval surveys. Collections were done in the domestic and peridomestic ecozones, and during rainy and dry seasons. Species composition, oviposition indices (ovitrap positive index (OPI), mean egg counts per ovitrap (MEO) and egg density index (EDI)) and *Stegomyia* indices (house index (HI), container index (CI) and Breteau index (BI)) were compared across the study areas, ecozones and seasons, and with the World Health Organization (WHO) DEN and YF epidemic thresholds.

**Results:** Overall, all the study areas were highly infested with diversified mosquito species, with the highest *Stegomyia* risk indices recorded in the urban areas. We identified (individuals: species) in the all study (70,550: 15), urban (32,331: 10), suburban (19,768: 14) and rural (18,451: 15) areas. Mosquitoes were mostly abundant in the domestic ecozones and rainy seasons. Larvae bred mainly in tires in the urban and suburban, and small containers in the rural areas. Wild *Aedes* species (*Aedes dendrophilus*, *Aedes lilii*, *Aedes fraseri*, *Aedes luteocephalus*, *Aedes metallicus* and *Aedes vittatus*) were mostly restricted in the suburban and rural areas, as well as mosquito predatory larvae (*Lutzia tigripes*, *Toxorhynchites* and *Eretmapodites*). However, *Aedes aegypti* spread and dominated throughout, showing highest proportions in the urban (90.2%), followed by the suburban (68.6%) and rural (68.5%) areas. OPI, MEO (egg/ovitrap/week) and EDI (egg/ovitrap/week) values were higher in the urban areas (53.0%, 8.7 and 16.4) than in the suburban (43.1%, 5.56 and 12.9) and rural (33.7%, 4.4 and 13.0) areas, respectively, and correlated with *Stegomyia* indices. HI, CI and BI were higher in the urban (48.0%, 40.4% and 61.8) compared with the suburban (37.8%, 32.0% and 43.3) and rural (34.5%, 28.87% and 39.8) areas. *Stegomyia* indices corresponded to the WHO-density scales of 6-8 in the urban and 5-7 in the suburban and rural areas. *Stegomyia* indices were all above the WHO epidemic thresholds: YF risk was high the urban and moderate in the suburban and rural areas, while people were permanently exposed to high risk of DEN outbreaks in all the areas.

**Conclusions/significance:** In Cocody-Bingerville, urbanization shifts mosquito biodiversity, with restriction of wild *Aedes* and predatory species in the rural and suburban areas and dominance of *Ae. aegypti* in the urban areas. *Stegomyia* indices exceeded the WHO DEN and YF epidemic thresholds in all the areas, with highest risks in the urban areas. This could explain the current DEN outbreaks in urban Cocody-Bingerville. Our study provides a baseline for raising local community awareness and guiding appropriate preventive actions, including managing identified larval habitats to contain ongoing dengue outbreaks in Côte d’Ivoire.

**Author summary:** Côte d’Ivoire has experienced several epidemics of dengue and yellow fever transmitted by *Aedes* mosquitoes. Over 80-90% of dengue and yellow fever cases were reported in the urban areas of the health district of Cocody-Bingerville, southeastern Côte d’Ivoire. The increase in the frequency, magnitude and burden of these outbreaks has paralleled to the fast and uncontrolled urbanization. However, data are still lacking on how urbanization influences the ecology of the local mosquito vectors and the risks of transmission of dengue and yellow fever, thus compromising adequate planning of preventive actions. Therefore, we assessed the mosquito species composition and larval breeding sites, and *Stegomyia* indices in urban, suburban and rural areas within the disease hotspots during ongoing dengue outbreaks in the Cocody-Bingerville. Our results showed that mosquito species diversity altered considerably from urban to rural areas, leading to a dominance of the main vector *Aedes aegypti* in all the study areas. Moreover, *Aedes aegypti* showed the highest proportions in the urban areas that contained more larval breeding sites (e.g., tires and small containers). *Stegomyia* indices were very high and above the World Health Organization (WHO) epidemic thresholds, suggesting that people were exposed to high dengue risks and moderate yellow fever risks in all the study areas, but were more exposed to both diseases in the urban areas. This could explain the recurrent and present dengue outbreaks in urban Cocody-Bingerville. A community-based management of identified larval habitats might help to control the ongoing dengue outbreaks in Cocody-Bingerville, and more widely in similar settings.

## Introduction

Many mosquito species in tropical and subtropical regions act as vectors of human and animal pathogens, including *Aedes*-borne arboviruses such as dengue virus (DENV), yellow fever virus (YFV), chikungunya virus (CHIKV), Zika virus (ZIKV), Rift Valley fever virus (RVFV) and West Nile virus (WNV), posing significant threats to the public and global health worldwide [1,2]. Arboviral epidemics are threatening over 831 million people (i.e., 70% of population) on the African continent [3]. The spread and geographical expansion of dengue (DEN), yellow fever (YF) and other arboviruses across African are mainly driven by urbanization and climate change [3–6]. Africa is projected to have the fastest urban growth rate in the world and will be home to an additional 950 million people, reaching 1.2 billion by 2050 [7]. In Africa, urbanization is mostly uncontrolled and unplanned, and thus provides millions of water-holding containers and people that can serve as ovipositing grounds and blood-feeding sources for the females of the main urban vector, *Aedes aegypti*, respectively [3,8–10], and can increase the transmission and burden of arboviruses [10–13]. In 2023 alone, 171,991 suspected cases of dengue, including 70,223 cases and 753 deaths were reported from 15 African countries, including Côte d’Ivoire [14]. Burkina Faso, a neighboring country of Côte d’Ivoire, only stands out as the most affected country with 146,878 suspected dengue cases and 688 deaths [15]. However, no licensed vaccines are still available for many arboviruses (except for yellow fever vaccine) and no well-structured routine programs are dedicated for *Aedes* vector control in most African countries, including Côte d’Ivoire. Therefore, the surveillance of *Aedes* vectors is crucial for the preparedness for and the response, prevention and control of arboviral epidemics.

The urban *Aedes aegypti* and *Aedes albopictus* species and many wild *Aedes* species (e.g., *Aedes metallicus*, *Aedes luteocephalus*, *Aedes vittatus*) colonize many urbanized seatings in Africa. These *Aedes* populations well-adapted to urban environments have highly anthropophilic behaviors and dwell within or in close proximity to human habitats (i.e., domestic and peridomestic premises) where females mostly blood-feed on humans and oviposit in vast arrays of man-made containers (e.g., tires, water receptables) [16]. Thus, urbanization can alter the ecology of *Aedes* species and the transmission of DENV and YFV. This major anthropogenic change can affect, at immature stages in breeding containers, the diversity and abundance of competitive [17,18], predatory (*Eretmapodites Lutzia*, *Toxorhynchites*) [19] and sympatric (*Culex* and *Anopheles*) species that can reduce or increase *Aedes* vector population size and density [16,20–22].

Côte d’Ivoire is one of most important emerging and re-emerging foci of DEN and YF and characterized by rapid urbanization in West Africa [23,24]. The urbanization rate raised from 32.0% in 1975 to 52.1% in 2021 [25,26]. The four serotypes of DENV (i.e., DENV1-4) and YFV are co-circulating in the country. From 2017 to 2024, Côte d’Ivoire has increasingly faced multiple of DEN outbreaks often coupled with YF cases. The majority (80-90%) of DEN and YF cases have occurred in the highly urbanized areas of the health district of Cocody-Bingerville (600,000 inhabitants), a part of the density capital city of Abidjan (7 million inhabitants) [27]. Cocody-Bingerville has faced multiple waves of DEN and YF epidemics in the recent years, 2017-2024 [23,24,28–31]. In 2017, DEN caused 623 suspected, 192 confirmed and 2 fatal cases [6].Out of the192 confirmed DEN cases, 66% were virus serotype 2 (DENV-2), 29% were DENV-3 and 5% were DENV-1 [32]. In 2019, outbreaks of DEN (3,201 suspected, 281 confirmed and 2 fatal cases) and YF (89 confirmed and 1 fatal cases) were reported [23,33]. In 2022, an DEN outbreak resulted in 181 suspected, 19 confirmed and 1 fatal cases [32] In 2023, DEN outbreaks have caused 321 infected cases and 27 deaths [34]. As of July 2024, 4,450 confirmed and 2 fatal DEN cases were recorded in Côte d’Ivoire, with the largest incidences recorded in Cocody-Bingerville [35].

Cocody-Bingerville is a suitable setting for monitoring the effects of increasing urbanization on the ecology of *Aedes* vectors and the risk of transmission of DENV and YFV. Indeed, the recent accelerations of arboviral outbreaks in Cocody-Bingerville have paralleled with a rapid and uncontrolled expansion of urbanizing fueled by the flourishing economic development and characterized by massive installations of people and a poor management of solid and plastic waste [36]. However, the effects of this important land-use change driven by fast urbanization on the egg-laying and larval ecologies of local mosquitoes and the DENV and YF outbreaks are still poorly understood, thus compromising adequate planning of sustainable prevention and control programs. Therefore, the present study aimed to determine the oviposition and larval ecologies of mosquitoes and DEN and YF epidemic risks along an urban-rural gradient during DEN outbreaks to better understand related underlying entomological mechanisms in Cocody-Bingerville. Since mosquito immatures are highly sensitive to human-mediated land-cover change, we hypothesized that urbanization would shape species composition and increase the density of *Ae. aegypti*, resulting in high DEN and YF transmission risks in urban areas. To validate this hypothesis, we monitored mosquitoes at all immature stages (eggs, larvae and pupae) and DEN and YF risks based on *Stegomyia* indices across urban, suburban and rural areas in Cocody-Bingerville using two highly sensitive standardized sampling methods, ovitraps (or oviposition traps) and larval surveys based on the “dipping” technique [37,38]. The outcomes provided critical information for both public health and research supporting the development of effective and sustainable *Aedes* vector control measures for preventing DEN, YF and other arboviral diseases in urban, suburban and rural settings of Cocody-Bingerville and Côte d’Ivoire, and more widely in African countries.

## Methods

### Study area

The study was carried out in the health district of Cocody-Bingerville (897,000 inhabitants). Cocody-Bingerville includes the municipalities of Cocody (5° 20’ 151 N; 3° 58’ W) and Bingerville (5° 21’ N; 3°54’ W) (Figure 1). Cocody-Bingerville is located in the region of Abidjan (7 million inhabitants), southeastern Côte d’Ivoire. Cocody-Bingerville is the main hotspots and foci of DEN and YF of Côte d’Ivoire. It accounts for 80-90% of DEN and YF cases reported in the country. From 2017 to 2024, Cocody-Bingerville has increasingly faced multiple resurgences of DEN and YF, with highest incidences and burden in urbanized areas [27,39,40]. *Ae. aegypti* densities, larval breeding sites (tires, discarded cans, water storage containers) and arboviral risks are high in the urban Cocody-Bingerville [16,20,39–41].

**Fig 1.**
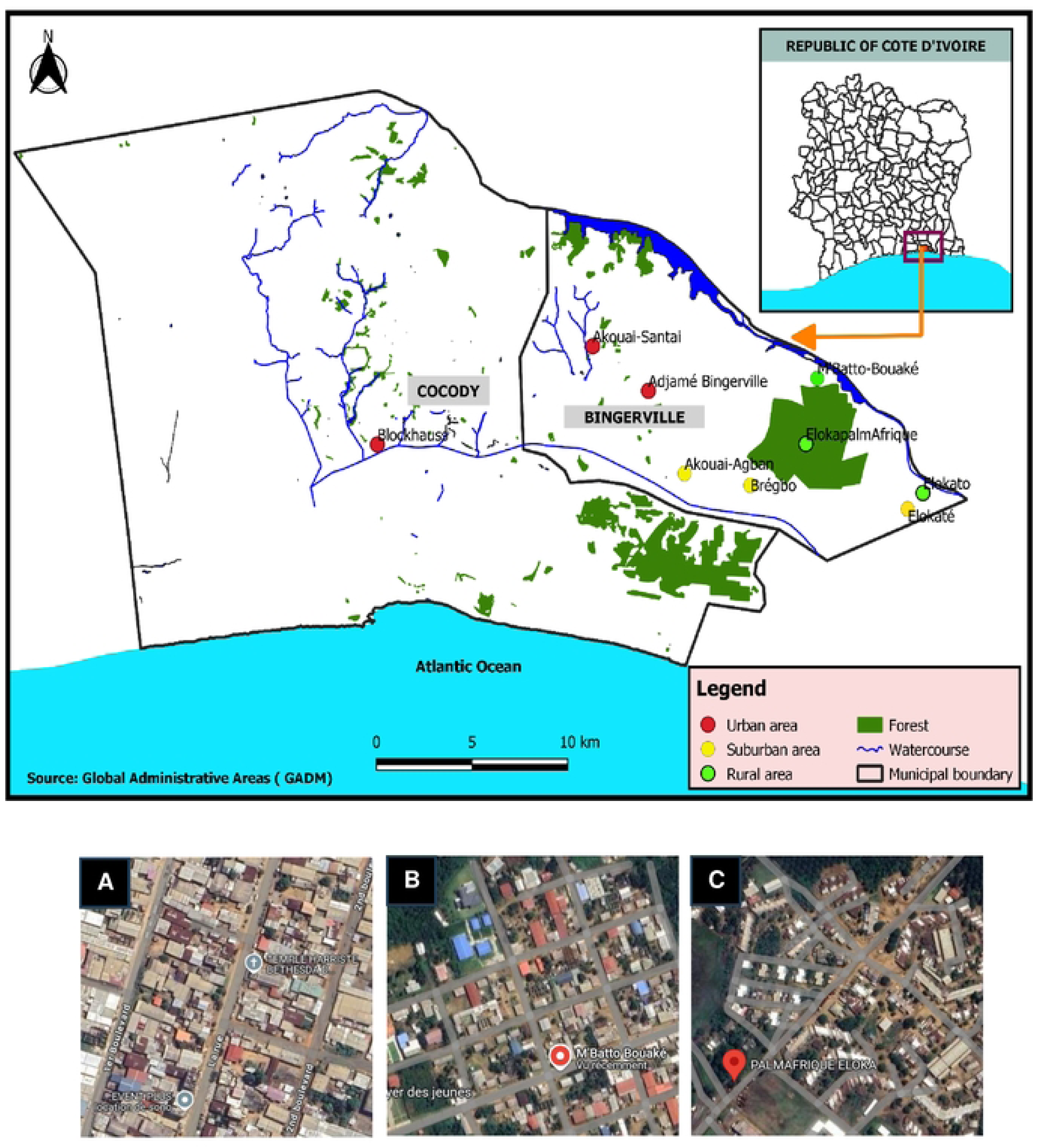
Locations of the study sites in alongside an urban-rural gradient in Cocody-Bingerville, southern Côte d’Ivoire. A: Urban areas (The urban areas include Adjamé-Bingerville, Blockhauss and Santé that are all urban neighbourhoods with numerous major and secondary paved roads and dense blocks of flats. These areas located in the highly urbanized parts of the municipalities of Cocody and Bingerville). B: Suburban areas (The suburban areas comprise the peri-urban towns of Akouai-Agban, Bregbo and Elokaté with only secondary paved roads and mixture of green spaces, agricultural areas and spaced houses). C: Rural areas (The rural areas cover the isolated rural villages of Elokato, Eloka-PalmAfrique and M’batto-Bouaké that the rural areas without major and secondary paved roads. These areas are close to the agricultural areas (oil palm and rubber farms), and forests). (PDF).

Cocody-Bingerville is subjected to accelerated and uncontrolled urbanization derived from the massive conversion of large and numbers of forests and farms (e.g., rubber and oil palm farms) into large settlements for people, resulting in three main land-cover types: urban, suburban and rural areas (Table 1). For our study, we randomly selected three sites per urbanization level: urban (Adjamé-Bingerville, Blockhauss and Santé), suburban (Akouai-Agban, Bregbo and Elokaté) and rural (Elokato, Eloka-PalmAfrique and M’batto-Bouaké). Only Blockhaus (5° 19’ 24” N, 4° 0’ 7” W) is located in the municipality of Cocody. Adjamé-Bingerville (5° 20’ 45” N, 3° 52’ 20” W), Santé (5° 21’ 56” N, 3° 53’ 55” W), Akouai-Agban (5° 18’ 39” N, 3° 51’ 20” W), Bregbo (5° 18’ 15” N, 3° 49’ 19” W) and Elokaté (5° 17’ 38” N, 3° 44’ 56” W), Elokato (5° 18’ 10” N, 3° 44’ 34” W), Eloka-PalmAfrique (5° 21’ 6” N, 3° 47’ 33” W) and M’batto-Bouaké (5° 19’ 0” N, 3° 48’ 2” W) are all located in the municipality of Bingerville. The urban areas are the urban neighbourhoods with numerous major and secondary paved roads and dense blocks of flats, located in the highly urbanized parts of Cocody and Bingerville. The total population in the three selected urban neighbourhoods is estimated at 9,200 inhabitants [25]. The suburban areas comprise the peri-urban towns with only secondary paved roads and mixture of green spaces, agricultural areas and spaced houses. The total population in the three selected suburban towns is estimated at 6,300 inhabitants [25]. The rural areas are the isolated rural villages without major and secondary paved roads. These areas are close to the agricultural areas (e.g., oil palm and rubber farms), and forests. The total population in the three selected villages is estimated at 4,900 inhabitants [25].

**Table 1.** Definition and classification of land-use and land-cover change and mosquito breeding sites sampled in arboviral hotspots in Cocody-Bingerville, southeastern Côte d’Ivoire. (PDF).

Cocody-Bingerville is located in a coastal area with tropical climate. The climate is hot and humid. There are four seasons distinguishable by the rainfall: long dry season (LDS) from December to March, long rainy season (LRS) from April to July, short dry season (SDS) from August to September and short rainy season (SRS) from October and November. Average annual precipitation ranges from 1,200 to 2,400 mm. The annual temperature is approximately 26.5 °C and the annual relative humidity ranges between 78 and 90%.

### Study design

This study was conducted along three urbanization levels: urban, suburban and rural (Table 1), with three equally sized study sites per level, totaling nine sites. Each study site was subdivided into two ecological zones (ecozones): domestic and peridomestic ecozones (Table 1). A cross-sectional survey was carried out in each of the four local climatic seasons (i.e., LRS, SRS, LDS and SDS) in each study site, resulting in four collection events per site and 36 collection events in total. Collections were done quarterly from August 2023 to July 2024.

### Ovitrap surveys

Standard WHO ovitraps were used to collect the eggs of *Aedes* mosquitoes [42,43]. Ovitraps were cut-out metal boxes with a volume of 400 cm^3^, garnished each with a 5 × 7 × 0.3 cm hardboard paddles, rough on one side and serving as an egg-laying substrate. Back-painted ovitraps were garnished with hardboard paddles and filled out at ¾ volume with distilled water mixed with rainwater (ratio: 1:1) [42] and left in the field for a one-week per each survey. Ovitraps were suspended 1.5 m above the ground and or installed at the ground level. The ovitrap devices were secured and protected from disturbances and rainfalls. The paddles, hatched larvae or pupae of mosquitoes, and remaining water from each ovitrap were collected and transferred separately into three labelled plastic cups. The label included the name of study site, collection point code, dates of ovitrap installation and collection. Each sampling point was georeferenced. We installed 100 ovitraps (50 in domestic ecozone and 50 in peridomestic ecozone) in each study site per survey. In total, we deployed 3,600 ovitraps (1,200 in urban, 1,200 in suburban and 1,200 in rural areas). The samples were transported in cool boxes to the laboratory for further processing.

### Larval surveys

In each study site, mosquito immatures (larvae and pupae) were collected from 100 randomly selected households. Collections were performed in domestic and peridomestic ecozones. Ready visible and accessible containers were inspected for the presence of mosquito larvae or pupae. When present, Immatures were sampled using flexible rubber tubing connected to a manual suction pump, ladles and pipettes depending on the size and shape of the breeding sites. Samples were stored separately in labelled plastic cups. Predatory mosquito larvae (*Lutzia tigripes*, *Eretmapodites* and *Toxorhynchites*) and other species (e.g. tadpoles) were immediately removed to avoid predation on other larvae. They were kept separately as described above. The plastic cups were labelled with information on household code, location, study site and date and transported to the insectarium. We recorded the ecozones and categories of inspected larval breeding sites. Breeding sites were classified into seven different categories: large containers, medium containers, small containers, tires, troughs, flowerpots and others according to the WHO guideline [44] (Table 1). The breeding sites sampled were toped-up at initial volume with mixture of distilled and rainwater (ratio: 1:1) in the domestic ecozones or rainwater only in the peridomestic ecozones. All the samples were transported in cool boxes to the field laboratory.

### Laboratory procedures

In the laboratory, the paddles were dried for seven days under ambient conditions (i.e., 25 ± 1 °C, 80-90% relative humidity and 12 hour:12 hour light:dark photoperiod). Paddles were covered with unimpregnated mosquito netting to prevent ovipositing from other mosquitoes. The paddles were then immersed separately in white-bottomed trays filled up to 75% capacity with distilled water for egg hatching. The drying-flooding process was repeated three times to maximize egg hatching.

Egg-derived and field-collected larvae were counted using a pipette and sorted by genus and stage. Larvae were reared to adults under similar ambient conditions as described above. In order to avoid overcrowding and limit mortality, a maximum of 20 larvae were placed in each 200-ml plastic cup filled at 75% capacity with distilled water and fed with Tetra-Min Baby Fish Food. Predatory larvae were fed with larvae from special colonies collected in the respective study sites. Emerged pupae collected were kept until adult emergence. Plastic cups were covered with fine mesh, unimpregnated mosquito netting fitted with sleeves. Each cup was labelled with the site, location and date. Emerged adults were identified morphologically under a binocular magnifying glass using available taxonomic keys [42,45]. Data was recorded in a designed database.

### Statistical analysis

Data were analyzed using Stata/MP 18.0. A significance level of 5% was set.

We calculated mosquito species richness (number of species) and abundance (number or proportion of individuals). We used proportion-test to compare mosquito proportions among, study areas, ecozones and seasons. We estimated other key biodiversity indicators such as species diversity, dominance and community similarity by Shannon-Weaver index (H), Simpson index (D) and Sorenson’s coefficient (CC), respectively [46,47]:

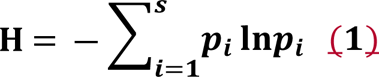

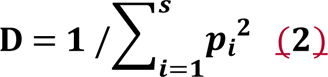

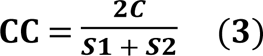

H: Shannon biodiversity index; *i*: a species in the study environment; *p_i_*: is the proportion of a species *i* to the total number of specimens found (N) in the study environment, calculated as follows: *p_i_* = n_i_/N where n_i_ is the number of specimens for the species; *S*: species richness; ln: natural logarithm; ∑: the sum of the calculations. *C* is the number of species that the two communities have in common, *S1* is the total number of species found in community 1, and *S2* is the total number of species found in community 2.

H is a statistical index of information that assumes that all species are represented in a sample and randomly sampled. The higher the value of H, the higher the species diversity; while the lower the value of H, the lower the species diversity. D is a dominance index, as it gives more weight to common or dominant species and assumes that a few rare species with only a few representatives will not affect diversity. The higher the D value, the higher the species abundance; whereas the lower the D value, the lower the species abundance. CC provides information on community similarity and helps to know how much two communities have overlap or similar. CC ranges from 0 to 1. The closer the value is to 1, the more the communities have species in common; complete community overlap is equal to 1; and complete community dissimilarity is equal to 0.

We compared species richness between the study areas, ecozones and seasons using one-way analysis of variance (ANOVA), followed by Bonferroni correction. The Shannon-Weaver species diversity index and Simpson dominance index were compared among the study areas, ecozones and seasons using Kruskal-Wallis test since the data exhibited significant deviations from normality (Shapiro-Wilk W test for normal data was significant, with p < 0.0001).

The numbers of *Aedes*-positive ovitraps and eggs were counted to calculate ovitrap positivity index (OPI), mean eggs count per ovitrap (MEO) and egg density index (EDI) [48]. OPI referred to the percentage of *Aedes*-positive ovitraps among total number of ovitraps examined. MEO was expressed as the mean number of eggs per number of ovitraps examined. EDI was defined as the mean number of eggs per *Aedes*-positive ovitraps. OPI, MEO and EDI were determined only for *Ae. aegypti* as this species represented 99.21% of *Aedes* genus. OPI, MEO and EDI were compared between study areas, ecozones and seasons using generalized linear mixed model (GLMM) procedures in order to consider the possible effects of and interactions between “study areas”, “study sites”, “ecozones” and “seasons’ and post-hoc tests of the trends [49].

The frequency of mosquito-positive breeding sites (FP) was expressed as the percentage of water-holding containers with at least one larva or pupa of mosquito (numerator) among the wet containers (denominator). The proportion of mosquito-positive breeding site types among the mosquito-positive breeding sites (PP) was estimated as the percentage of each mosquito-positive container type (numerator) among the total mosquito-positive containers (denominator) in each study area. Chi-square test (χ^2^) was used to compare the proportions of mosquito-positive breeding sites between the study areas, ecozones and seasons.

DENV and YFV transmission risk was determined using *Ae. aegypti* larval indices, termed *Stegomyia* indices, that included container index (CI: percentage of *Ae. aegypti*-positive containers among containers examined), house index (HI: percentage of houses containing at least one *Ae. aegypti*-positive breeding site) and Breteau index (BI: number of *Ae. aegypti*-positive containers per 100 houses inspected). A container is considered positive when it contains at least one larva or pupa of *Ae. aegypti*. A house is positive when it contains at least one *Ae. aegypti*-positive container. HI, CI and BI were compared across the study areas and seasons using one-way ANOVA. We used HI, CI and BI values to estimate the levels of DEN and YF outbreak risks according to the WHO epidemic thresholds, as follows:

- For DEN: If CI > 3% or HI > 4% and BI > 5, epidemic risk is high [50].
- For YF: If IC < 3%, epidemic risk is low. 3% ≤ CI ≤ 20%: risk is moderate, CI > 20%, epidemic risk is high; If HI < 4%, epidemic risk is low. 4% ≤ HI ≤ 35%: epidemic risk is moderate. HI > 35%: epidemic risk is high; and If BI < 5, epidemic risk is low. 5 ≤ BI ≤ 50: risk is moderate. BI > 50: epidemic risk is high [51].

We tested correlations between the ovitrap indices (OPI, MEO and EDI) and *Stegomyia* indices (CI, HI and BI) using Pearson’ s test. We used Spearman’s test only for any tests involving OPI as OPI data were not normally distributed (Shapiro-Wilk W test was significant, p < 0.05).

### Ethics statement

Before sample collection, the study protocol received ethical approval from the national ethics committee of Côte d’Ivoire (N/Ref: 071-23/MSHPCMU/CNESVS-km). Additionally, permissions were obtained for the General Direction of Health (GDH) of the Ministry of Health (MoH) of Côte d’Ivoire, the administrative and health authorities of Cocody-Bingerville and the local community leaders of the study areas. Mosquito samples and data were collected with the authorization of the residents and/or owners. This study did not involve endangered or protected species.

## Results

### Species composition

A cumulative total of 70,550 mosquitoes (33,683 females and 36,867 males) was identified in the current study (Table 2). The fauna comprised several species of public health and ecological interests, including disease vector, predatory and sympatric species. The collected mosquito fauna belonged to six genera, dominated by *Aedes* (79.10%, n = 55,804), followed by *Culex* (17.63%, n = 12,436), *Lutzia* (1.76%, n = 1,239), *Eretmapodites* (0.93%, n = 653), *Anopheles* (0.58%, n = 411) and *Toxorhynchites* (0.01%, n = 7). We identified 15 species among which *Ae. aegypti* was the most abundant accounting for 78.48% (n = 55,366).

**Table 2.** Species composition of adult mosquitoes emerged from eggs and larvae collected along an urban-rural gradient in the arboviral hotspots of Cocody-Bingerville, southeastern Côte d’Ivoire from August 2023 to July 2024. (PDF).

Species composition varied substantially between the study areas. The highest number of specimens (n = 32,331 individuals) was found in the urban areas, while the highest species richness (14 species) was recorded in the rural areas. Similarly, *Aedes* arbovirus vectors showed the highest proportion and lowest species richness in the urban (90.42% and 3 species) and lowest proportion and highest species richness in rural areas (69.45% and 7 species). *Aedes aegypti* was mostly abundant in the urban (90.22%, 29,168/32,330), followed by the suburban (68.60%, 13,560/19,768) and rural (168.49%, 2,638/18,451) areas. *Aedes vittatus* had low proportions but was present in all the study areas. Other potential arbovirus vectors, including *Aedes* species (*Aedes dendrophilus*, *Aedes lilii*, *Aedes luteocephalus* and *Aedes metallicus*), *Culex* species (*Culex quinquefasciatus*) and *Eretmapodites* species (*Eretmapodites chrysogaster* and *Eretmapodites quinquevittatus*) were mostly collected in the rural and suburban areas.

Immature mosquitoes able to influence the ecology of *Aedes* and the transmission of DENV and YFV were sampled in the study areas. The sympatric *Culex* species (*Cx. quinquefasciatus* and *Culex nebulosus*) had proportions above 7% in the urban, and represented 22% of the total fauna in the suburban and rural areas. Mosquito predatory larvae (*Eretmapodites*, *Lutzia* and *Toxorhynchites*) were mostly found in the suburban and rural areas. Small proportions of the malaria mosquito *Anopheles gambiae* was collected from containers in the all the study areas.

### Biodiversity

The mean species richness values were 5.17 (95% CI: [4.59, 5.75]) in the urban, 5.33 (95% CI: [4.54, 6.13]) in the suburban and 5.50 (95% CI: [4.80, 6.20]) in the rural areas (Table 3). The species richness was statistically comparable between the study areas (F = 0.24, df = 2, p = 0.7843). The respective mean values of Shannon diversity index were estimated at 0.43 (95% CI: [0.30, 0.56]), 0.86 (95% CI: [0.73, 0.99]) and 0.86 (95% CI: [0.73, 0.99]) in the urban, suburban and rural areas and with statistical differences between the study areas (χ^2^ = 21.766, df = 2, p < 0.0001). The Shannon diversity index values were significantly lower in the urban areas compared with the suburban areas (χ^2^ = 15.349, df = 1, p < 0.0001), and the rural areas (χ^2^ = 17.092, df = 1, p < 0.0001), but did not differ statistically between suburban in the rural areas (F = 0.002, df = 1, p = 0.9812). The Simpson dominance index was 0.20 (95% CI: [0.13, 0.28]) in the urban areas, 0.46 (95% CI: [0.37, 0.55]) in the suburban areas and 0.45 (95% CI: [0.38, 0.52]) in the rural areas. Simpson indices significantly differ between the study areas (χ^2^ = 21.880, df = 2, p < 0.0001). Simpson index was significantly lower in the urban areas than in the suburban areas (χ^2^ = 15.027, df = 1, p < 0.0001) and in the rural areas (χ^2^ = 17.607, df = 1, p < 0.0001). No significant difference was detected in the Simpson index between the suburban and rural areas (F = 0.01, df = 1, p = 0.822). Considering the urban areas as the reference (CC = 1), the Sorenson coefficients were close to 1 in both the suburban areas (CC = 0.83) and the rural areas (CC = 0.80), revealing that the three study areas shared at least 80% of species (Table 3).

**Table 3.** Biodiversity of adult mosquitoes collected emerged from eggs and larvae sampled across an urban-rural gradient in the arboviral hotspots of Cocody-Bingerville, southeastern Côte d’Ivoire from August 2023 to July 2024. (PDF).

### Geographical and seasonal variations

Overall, mosquito species was more diversified in the domestic ecozones (14 species) compared with the peridomestic ecozones (12 species) (S1 Table). The species richness was equal between domestic and peridomestic ecozones in the urban areas (9 species) and suburban areas (11 species), but higher in the domestic ecozones (12 species) compared with peridomestic ecozones (8 species) in the rural areas. Overall, the proportions of mosquitoes were significantly higher in the domestic ecozones (57.56%, 40,608/70,550) than in the peridomestic ecozones (42.44%, 29,942/70,550) (χ^2^ = 3224.4, df = 1, p < 0.0001). *Aedes aegypti* followed the same distributional patterns, displaying statistically higher propositions in the domestic ecozones compared to the peridomestic ecozones (domestic *vs* peridomestic) in all the study areas (54.72% *vs* 45.28%; χ^2^ = 987.70, df = 1, p < 0.0001), the suburban areas (60.86% *vs* 39.14%; χ^2^ = 1279.20, df = 1, p < 0.0001) and rural areas (59.16% *vs* 40.84%; χ^2^ = 848.11, df = 1, p < 0.0001). However, *Ae. aegypti* proportions were statistically comparable between the domestic and peridomestic ecozones in the urban areas (49.95% *vs* 50.05; χ^2^ = 0.06, df = 1, p = 0.7974). The other *Aedes* species were mostly collected in the domestic ecozones.

The proportions of mosquitoes differed significantly over the seasons (χ^2^= 12158, df = 3, p < 0.0001) (Figure 2A). Proportions showed the highest values during LDS, and the lowest values during SRS in all the study areas and in the domestic and peridomestic ecozones. Similarly, *Ae. aegypti* exhibited the highest proportions in LDS and the lowest proportions in SRS in all the study areas and the domestic and peridomestic ecozones, as well. Compared with peridomestic ecozones during all seasons, *Ae. aegypti* showed the significantly highest proportions in the domestic ecozones during SRS in urban (52.73% *vs* 47.27%, χ^2^ = 31.095, df = 1, p < 0.0001), SDS in suburban (71.19% *vs* 28.81, χ^2^ = 623, df = 1, p < 0.0001) and LDS in rural (65.71% *vs* 34.29%; χ^2^ = 381.96, df = 1, p < 0.0001) areas (Figure 2B). *Aedes aegypti* presented statistically similar proportions among domestic and peridomestic ecozones across the seasons (p > 0.05).

**Fig 2.**
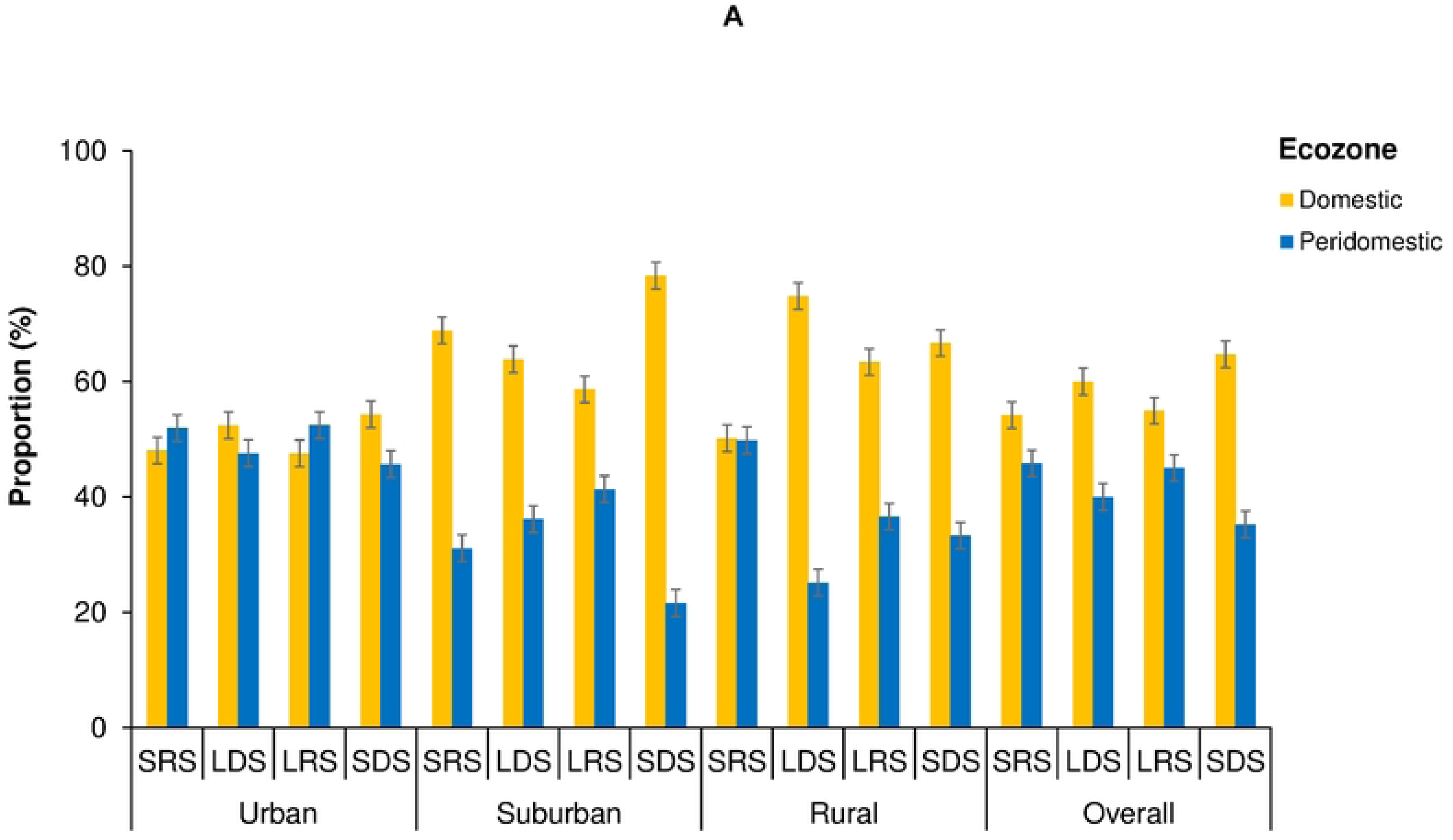
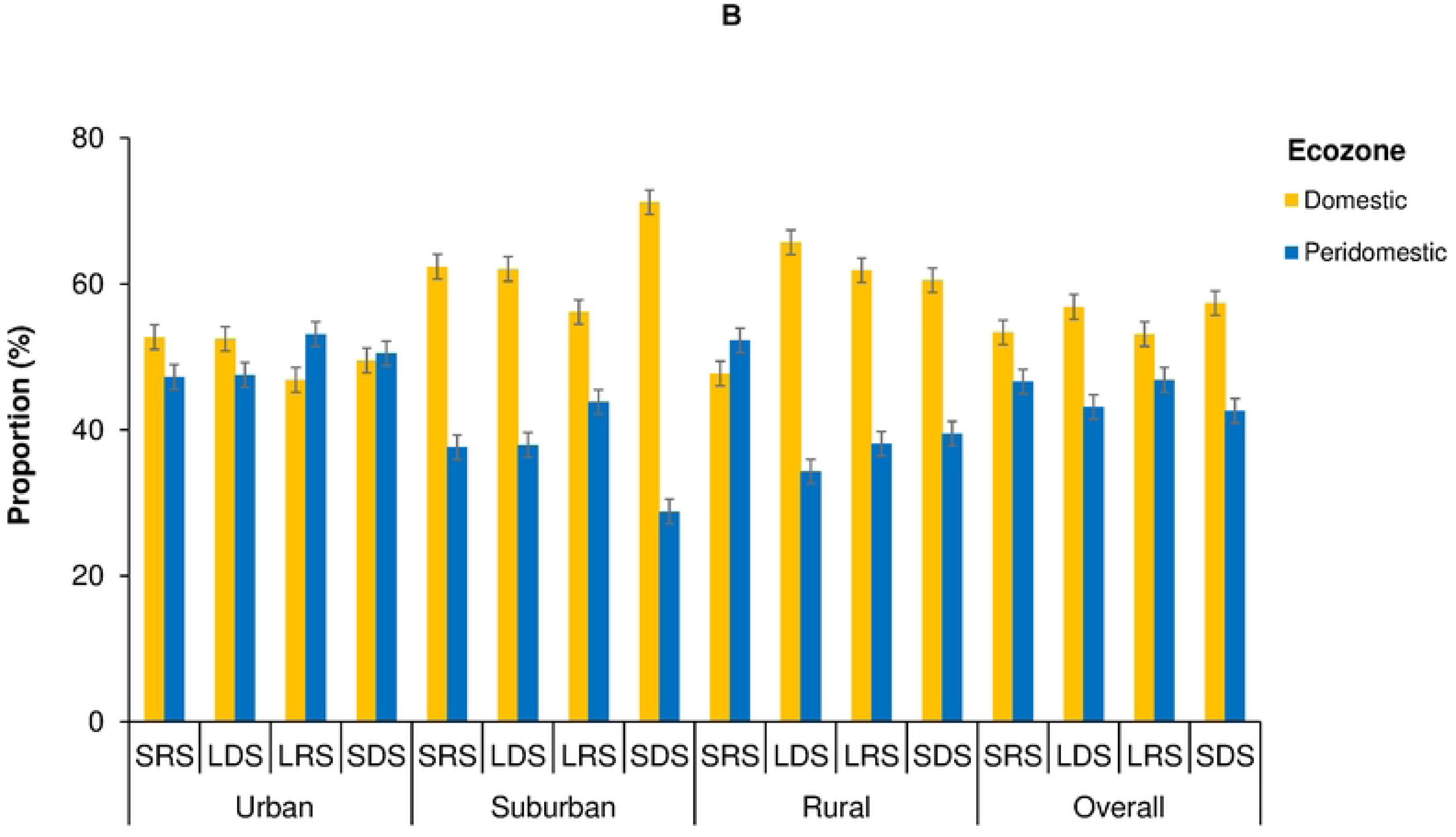
Seasonal variations of the proportions of adult mosquitoes emerged from eggs and larvae collected in the ecozones along an urban-rural gradient in Cocody-Bingerville, southeastern Côte d’Ivoire from August 2023 to July 2024. A: Mosquito fauna, B: *Aedes aegypti*. SRS: short rainy season, LDS: long dry season, LRS: long rainy season, SDS: short dry season. Error bars show the confidence intervals (95% CI). (PDF).

### Oviposition ecology

#### Species composition of egg-derived adults

In total, 21,521 adult mosquitoes (10,578 females and 10,943 males) belonging to nine species derived from collected eggs (Table 2). The respective totals of 9,382 (three genera and six species), 6,468 (four genera and seven species) and 5,671 (four genera and eight species) mosquitoes were found in the urban, suburban and rural areas. Taken together, the proportions of the egg-derived mosquitoes were significantly higher in the urban areas (43.57%, 9,382/21,521), compared to the suburban areas (27.50%, 6,468/21,521) and rural areas (28.93%, 5,671/21,521) (χ^2^ = 1534, df = 2, p < 0.0001). *Aedes aegypti* was the most prevalent species in all the study areas, with higher proportion in the urban areas (98.83%, 9,272/9,382), followed by suburban areas (95.16%, 6,155/6,177) and the rural areas (90.13%, 5,111/5,116).

#### Ovitrap indices of *Aedes aegypti*

##### Ovitrap positive index

Table 4 displays the variations of OPI across the study areas. OPI value was of 42.99% (95% CI: [41.31%, 44.67%]) for all the study areas. OPI was higher in the urban (53.04%, 95% CI: [50.04%-56.04%]), followed by the suburban (43.05%, 95% CI: [40.13%, 45.97%]) and rural (33.73%, 95% CI: [31.02%, 36.45%]) areas. GLMM showed that OPI differed significantly between the urban and suburban areas (Coefficient = −20.70, Z = −2.06, p < 0.0001). However, OPI values were statistically similar between the urban and rural areas (Coefficient = −4.47, Z = −0.46, p = 0.647), between the suburban and rural areas (Coefficient = 16.23, Z = 1.61, p = 0.107).

**Table 4.** *Aedes aegypti* oviposition indices across an urban, rural gradient in the arboviral hotspots of Cocody, Bingerville, southeastern Côte d’Ivoire from August 2023 to July 2024. (PDF).

OPI values in the domestic and peridomestic ecozones were estimated at 57.79% (95% CI: [53.66%, 61.92%]) and 47.97% (95% CI: [43.65%, 52.29%]) in the urban, 42.58% (95% CI: [38.49%, 46.67%]) and 43.05% (95% CI: [40.13%, 45.97%]) in the suburban, and 32.25% (95% CI: [28.44%, 36.05%]) and 35.21% (95% CI: [31.33%, 39.10%]) in the rural areas, respectively. OPI values were significantly higher in the domestic ecozones than in the peridomestic ecozones in the urban areas (contrast = −9.04, Z = −3.17, p = 0.002), but comparable among both ecozones in the suburban (contrast = 1.24, Z = 0.44, p = 0.657) and rural (contrast = 2.76, Z = 1.02, p = 0.309) areas. The highest and lowest values of OPI were recorded during LDS (64.71%, 95% CI: [58.80%, 70.61%]) and SRS (42.20%, 95% CI: [36.40%, 48.00%]) in the urban, LDS (61.70%, 95% CI: [55.99%, 67.41%]) and SRS (22.00%, 95% CI: [17.29%, 26.71%]) in the suburban, and LRS (41.98%, 95% CI: [36.30%, 47.66%]) and SRS (28.09%, 95% CI: [22.97%, 33.22%]) in the rural areas, respectively. These highest and lowest values were significantly different in the urban (contrast = −21.48, Z = −5.31, p < 0.0001), suburban (contrast = −39.46813, Z = −10.27, p < 0.0001) and rural (contrast = −14.05809, Z = −3.69, p < 0.0001) areas. OPI varied among the domestic and peridomestic ecozones over seasonal variations (S2 Table). The effects and interactions among the study areas, sites, ecozones and seasons, and trends (linear, quadratic and cubic trends) in OPI values over the seasons were statistically significant (Table 5 and S3 Table).

**Table 5.** Effects, interactions and trends in *Aedes aegypti* oviposition indices along an urban-rural gradient in the arboviral hotspots of Cocody-Bingerville, southeastern Côte d’Ivoire from August 2023 to July 2024. (PDF).

##### Mean egg counts per ovitrap

MEO was estimated at 6.14 (95% CI: (5.75-6.53]) egg/ovitrap/week for all the study areas (Table 4). MEO showed the highest values in the urban (8.67, 95% CI: [7.89, 9.46]) egg/ovitrap/week), followed by the suburban (5.56, 95% CI: [4.95, 6.16]) egg/ovitrap/week) and the rural (4.38, 95% CI: [3.78, 4.97]) egg/ovitrap/week) areas. MEO values were significantly different between the urban and suburban areas (Coefficient = −5.84, Z = −2.48, p < 0.0001). MEO values were statistically similar between the urban and rural areas (Coefficient = −4.47, Z = −1.36, p = 0.175), and between the suburban and rural areas (Coefficient = 2.74, Z = 1.16, p = 0.245).

The respective values of MEO in the domestic and peridomestic ecozones were 9.32 (95% CI: (8.22, 10.42]) and 7.98 (95% CI:[ 6.86, 9.10]) egg/ovitrap/week in the urban, 5.46 (95% CI: [4.63, 6.31]) and 5.64 (95% CI: [4.77-6.52]) egg/ovitrap/week in the suburban, and 3.71 (95% CI: [3.00, 4.43]) and 5.03 (95% CI: [4.09, 5.98]) egg/ovitrap/week in the rural areas. MEO values were statistically similar between the domestic and peridomestic ecozones in the urban areas (contrast = −-1.01, Z = −1.52, p = 0.130), in the suburban (contrast = 0.16, Z = 0.25, p = 0.799) and rural (contrast = 1.181462, Z = 1.86, p = 0.062) areas. MEO showed the highest and lowest values during LRS (10.69 (95% CI: [8.78, 12.61]) egg/ovitrap/week) and SRS (6.30 (95% CI: [5.09, 7.51%])) egg/ovitrap/week in the urban, LDS (8.02 (95% CI: [6.72, 9.31]) egg/ovitrap/week) and SRS (3.17 (95% CI: [2.22, 4.11]) egg/ovitrap/week) in the suburban, and LRS (7.53 (95% CI: [5.79, 9.28]) egg/ovitrap/week) and LDS (2.33 (95% CI: [1.76, 2.90]) egg/ovitrap/week) in the rural areas, respectively. These highest and lowest values of MEO differed significantly in the urban (contrast = −4.39, Z = −4.71, p < 0.0001), suburban (contrast = −4.76, Z = −5.29, p < 0.0001) and rural (contrast = −5.19, Z = −5.73, p < 0.0001) areas. MEO values altered across the domestic and peridomestic ecozones according to the seasons (S2 Table). The effects and interactions among the study areas, sites, ecozones and seasons, and trends (linear, quadratic and cubic trends) in MEO across the seasonality were statistically significant (Table 5 and S4 Table).

##### Egg density index

EDI was estimated at 14.28 (95% CI: [13.58, 14.99]) egg/ovitrap/week for all the study areas (Table 4). The urban areas had the highest EDI values (16.35, 95% CI: [15.19, 17.51]) egg/ovitrap/week), followed by the rural areas (12.97, 95% CI: [11.56, 14.39]) egg/ovitrap/week) and the suburban areas (12.90, 95% CI: [11.80, 14.00]) egg/ovitrap/week). No significant differences were detected in EDI values between the urban and suburban areas (Coefficient = −7.27, Z = −1.67, p = 0.094), and between the urban and rural areas (Coefficient = −4.67, Z = −1.24, p = 0.214). However, EDI was statistically different between the suburban and rural areas (Coefficient = 11.53, Z = 2.57, P = 0.010).

EDI values in the domestic and peri-domestic ecozones were 16.13 (95% CI: [14.61, 17.64]) and 16.64 (95% CI: [14.84, 18.44]) egg/ovitrap/week in the urban, 12.84 (95% CI: [11.29, 14.39]) and 12.96 (95% CI: [11.38, 6.52]) egg/ovitrap/week in the suburban, and 11.51 (95% CI: [9.75, 13.26]) and 14.31 (95% CI: [12.12, 16.48]) egg/ovitrap/week in the rural areas. EDI values were statistically comparable between the domestic and peri-domestic ecozones in the urban areas (contrast = 0.68, Z = 0.60, p = 0.548), in the suburban (contrast = −0.11, Z = −0.09, p = 0.930) and rural (contrast = 1.69, Z = 1.11, p = 0.265) areas. EDI showed the highest and lowest values during LRS (18.97, 95% CI: [16.24, 21.71]) egg/ovitrap/week) and LDS (13.30 (95% CI: [11.45, 15.15]) egg/ovitrap/week) in the urban, SRS (14.40, 95% CI: [11.36, 17.46]) egg/ovitrap/week) and SDS (12.16 (95% CI: [9.09, 15.25]) egg/ovitrap/week) in the suburban, and LRS (17.96, 95% CI: [14.56, 21.36]) egg/ovitrap/week) and LDS (6.95, 95% CI: [5.68, 8.21]) egg/ovitrap/week) in the rural areas, respectively. These highest and lowest values were significantly different in the urban (contrast = −6.07, Z = −4.00, p < 0.0001), the suburban (contrast = 3.78, Z = 1.65, p = 0.099) and rural (contrast = −8.64, Z = −4.09, p < 0.0001) areas. EDI values changed among the domestic and peridomestic ecozones across the seasonality (S2 Table). The effects and interactions among the study areas, sites, ecozones and seasons, and trends (linear, quadratic and cubic trends) in EDI across the seasonal variability were almost statistically significant (Table 5 and S5 Table).

### Larval ecology

#### Species composition of larvae-derived adults

A total of 49,029 adult mosquitoes (23,106 females and 25,924 males) emerged from larvae and pupae sampled in all the study areas (Table 2). The species richness was lower in the urban areas (9 species), compared with the suburban areas (13 species) and the rural areas (14 species). Conversely, the urban areas had the highest number of specimens (n = 22,949), followed by the suburban areas (n = 13,300) and the rural areas (n = 12,780). *Aedes* genus showed the greatest proportions in all the study areas (71.87%, 35,238/49,029), exhibiting strong proportions that decreased from 86.97% (19,959/22,949) in the urban areas to 60.24% (7,699/12,780) in the rural areas and 57.00% (7,580/13,300) in the suburban areas (Table 2). *Aedes* species richness increased from the urban areas (3 species) to the suburban areas (5 species) and rural areas (7 species). *Aedes aegypti* dominated the mosquito fauna, showing respective proportions of 86.70% (19,896/22,949) in the urban areas, 55.68% (7,405/13,300) in the suburban areas and 58.90% (7,527/12,780) in the rural areas. Predatory larvae of *Er. chrysogaster, Er. quinquevittatus* and *Lu. tigripes* were sampled in all study areas, while *Tx. brevipalpis* was confined to the suburban and rural areas. Unexpected non-container specialist larvae of mosquitoes, such as *An. gambiae*, were sampled from containers in all the study areas, but at low proportions of 1.05% (240/22,949) in the urban, 0.40% (n = 53/13,300) in the suburban and 0.92% (118/12,780) in the rural areas.

#### Larval breeding sites

A total of 5,054 potential larval breeding containers were examined in all study areas (S6 Table). The urban areas (FP= 66.1%, 1,211/1,833) had the highest proportions of mosquito-positive breeding sites, followed by the suburban areas (FP = 65.8%, 1,069/1,624) and rural areas (63.2%, 1,010/1,567) (Fig 3A). Overall, the mosquito-positive breeding sites were ranked by proportions in descending order, as follows: tires (FP = 74.28%, 1,363/1,835), flowerpots (FP = 70%, 42/60), other types of containers (FP = 68,97%, 289/419), small containers (FP = 65,90%, 887/1346), drinking troughs (FP = 56.56%, 69/122), medium containers (FP = 55.75%, 441/791) and large containers (FP = 41.37%, 199/481). In all the study areas, most of the positive breeding sites were colonized by the larvae or pupae of *Aedes* species, mostly identified as *Ae. aegypti*.

**Fig. 3.**
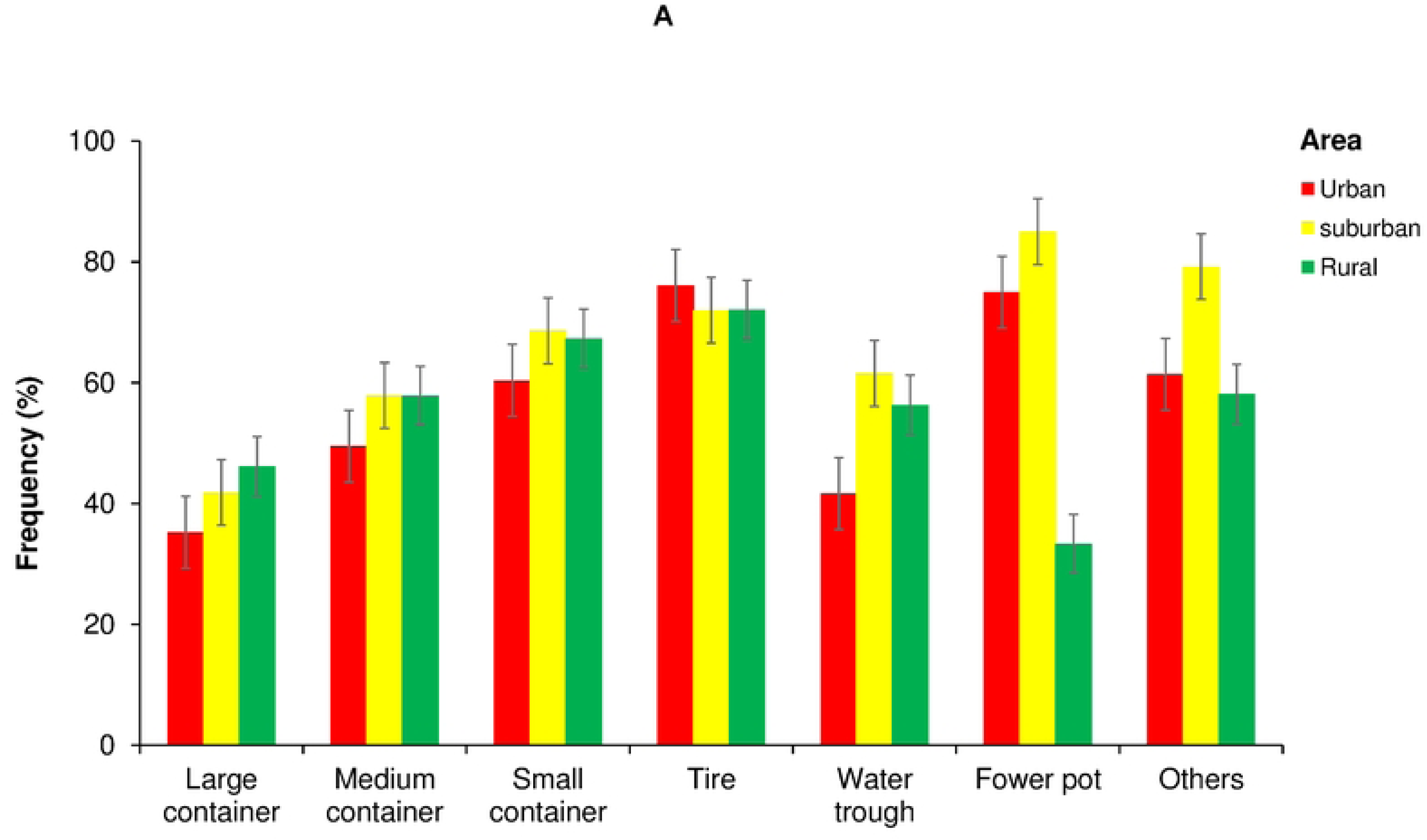
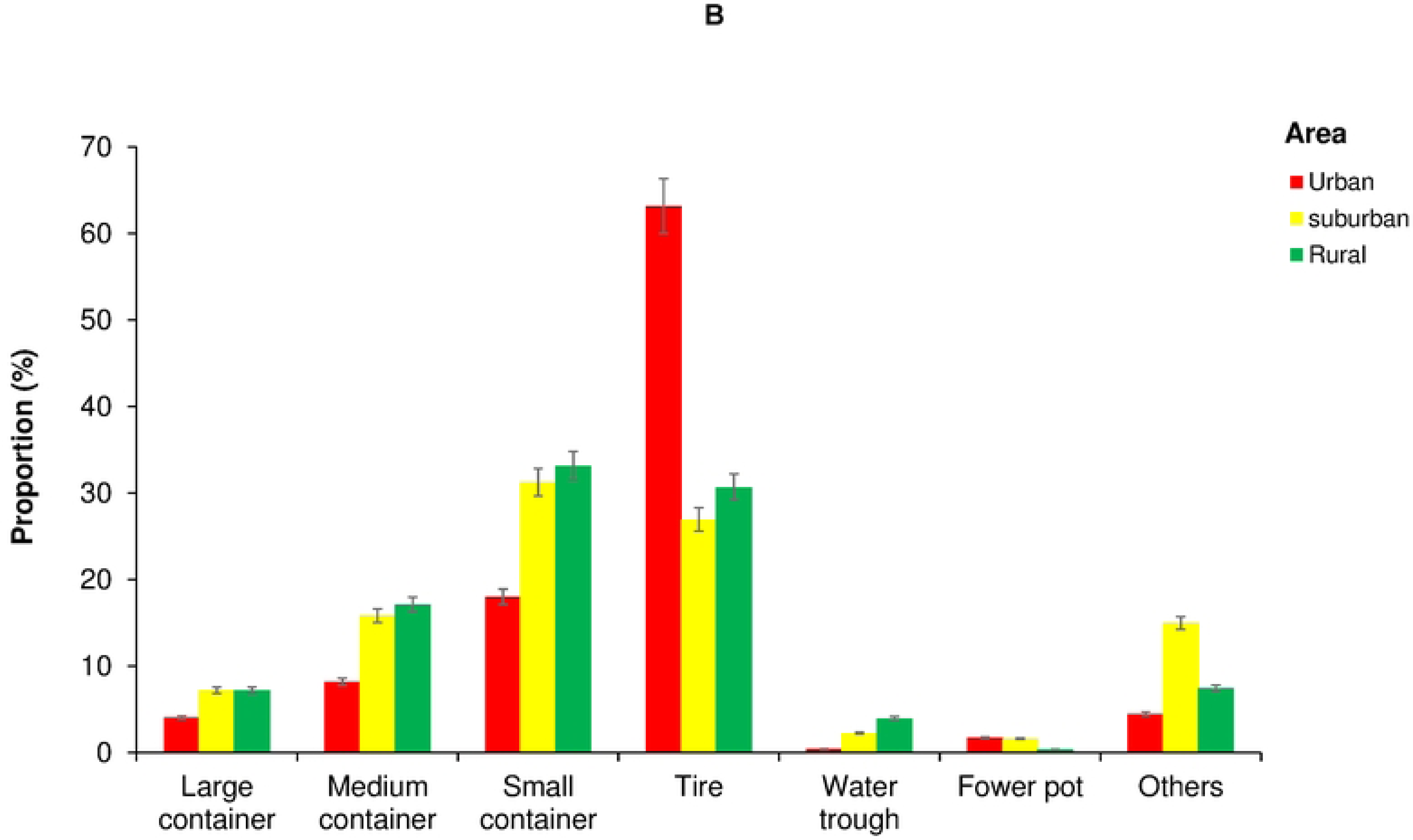
Dynamics of mosquito breeding sites in across an urban-rural gradient in Cocody-Bingerville, southeastern Côte d’Ivoire from August 2023 to July 2024. Error bars show the confidence intervals (95% CI). (PDF).

Of the 3,290 (65.10%) mosquito-positive breeding sites, the most were tires (PP = 41.43%, 1,363/3,290), followed by small containers (PP = 26.96%, 887/3,290), medium containers (PP = 13.40%, 441/3,290), other types (PP = 8.78%, 289/3,290), large containers (PP = 6.05%, 199/3,290), drinking troughs (PP = 2.10%, 69/3,290) and flowerpots (PP = 1.28%, 42/3,290) (Fig 3B). The proportions of these mosquito-positive sites were significantly higher in the urban areas (36.81%, 1,211/3,290) than in the suburban areas (32.49%, 1,069/3,290) and the rural areas (30.69%, 1,010/3,290) (χ^2^ = 29.2, df = 2, p < 0.0001). The urban areas, tires were most positive containers (PP = 63.17%, 765/1,211), followed by small containers (PP = 18%, 218/1211). Small containers were more common in the suburban (PP = 31.24%, 334/1,069) and rural (PP = 33.17%, 335/1010) areas. Additionally, tires displayed strong positivity in the suburban (26,94%, 288/1,069) and rural (30.69%, 310/1,010) areas (Fig 3B).

#### Geographical and seasonal distribution of the larval breeding sites

The domestic ecozones had significantly the highest number of mosquito-positive breeding sites compared with the peridomestic zones (χ^2^ = 623.81, df = 1, p < 0.0001) (Fig 4A). The domestic ecozones accounted for 52.52% (636/1,211) of mosquito-positive breeding sites in the urban areas, 76.89% (822/1,069) in the suburban areas and 68.71% (694/1,010) in the rural areas. Tires and small containers were the most abundant positive in all the two domestic and peridomestic ecozones. Additionally, the highest proportions of mosquito-positive breeding sites were recorded in SRS (FP = 70.7%, 403/554) in the urban areas, LRS (FP = 69.3%, 345/ 498) in the suburban areas and SDS (70.3%, 185/263) in the rural areas (S6 Table). *Aedes aegypti* the breeding site positivity showed the highest rates during LRS in the urban (33,60%, 249/741), suburban (32,31%, 168/520) and rural (34,59%, 165/477) areas (Fig 4B).

**Fig. 4.**
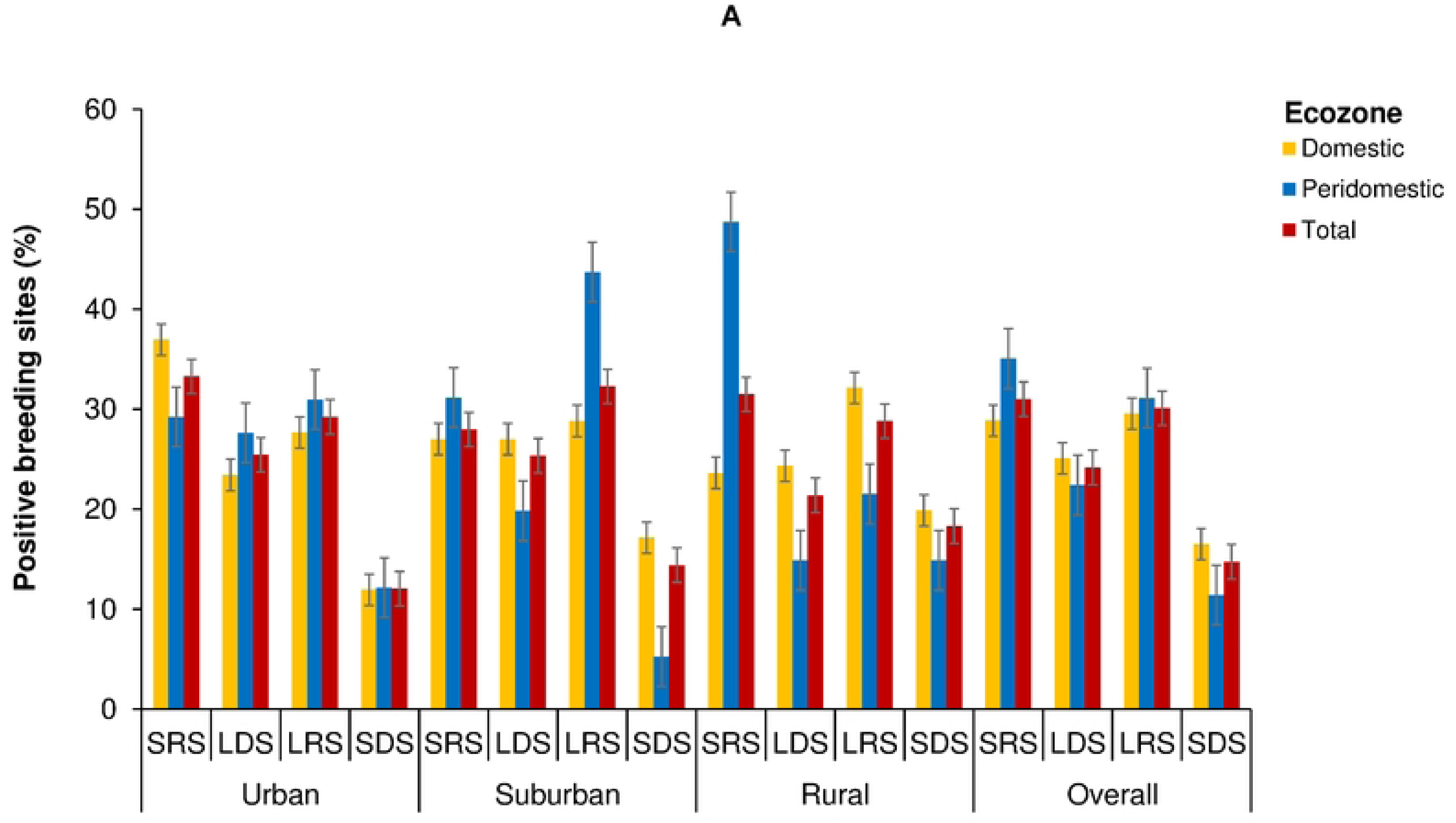
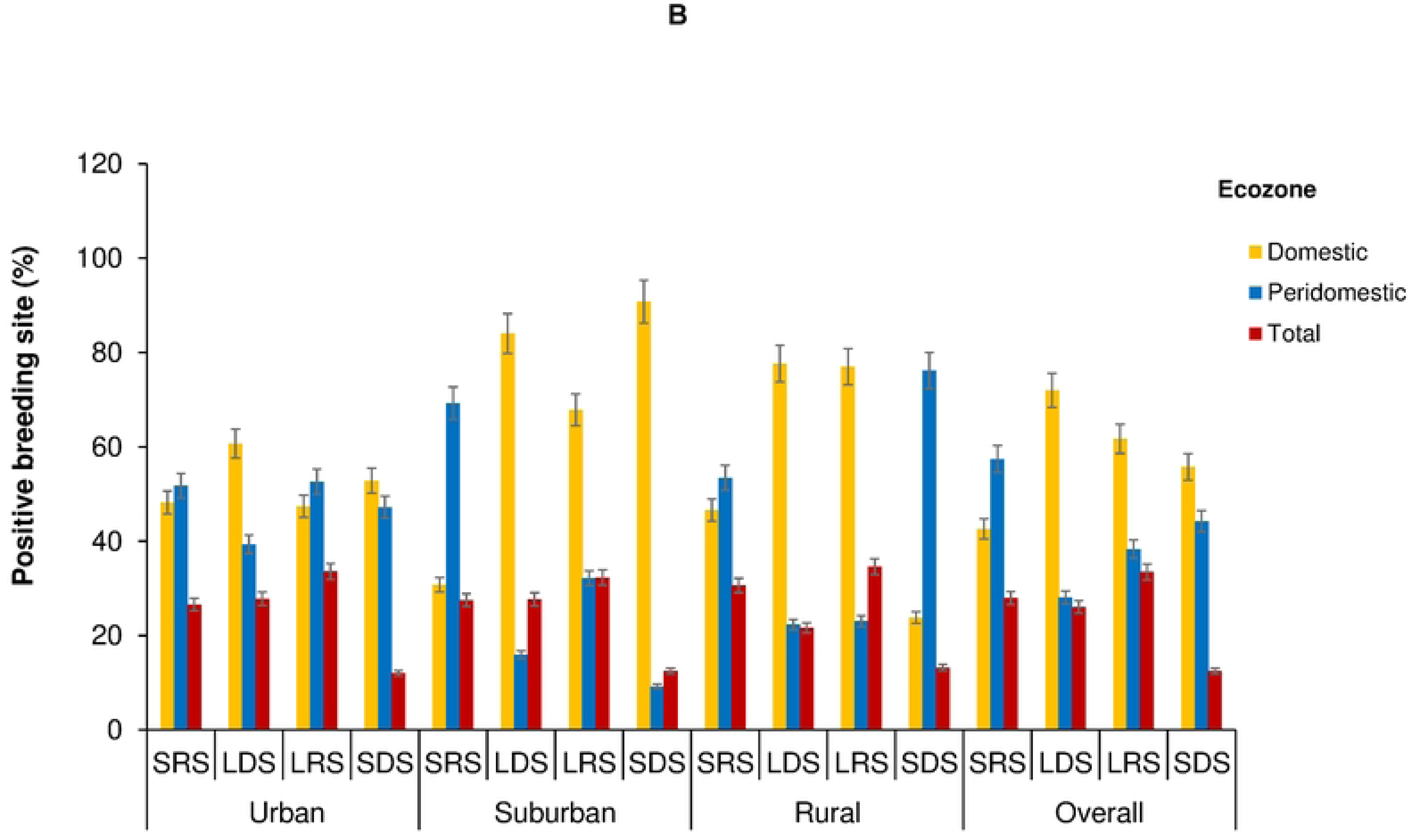
Geographical distribution of breeding sites mosquitoes (A) and *Aedes aegypti* (B) in across an urban-rural gradient in Cocody-Bingerville, southeastern Côte d’Ivoire from August 2023 to July 2024. A: Mosquito fauna, B: *Aedes aegypti*. SRS: short rainy season, LDS: long dry season, LRS: long rainy season, SDS: short dry season. Error bars show the confidence intervals (95% CI). (PDF).

#### Container productivity

The species richness and abundance of mosquito immatures varied among various water-holding containers sampled in the urban, suburban and rural areas (S7 Table). Immatures were mostly collected in tires (46.16%, 22,632/49,029) and small containers (24.56%, 22,632/49,029) all the study areas. Tires harbored the highest proportions of immatures in the urban areas (65.99%, 15,143/22,949), followed by the suburban (30.12%, 4,403/14,617) and rural (26.92, 3086/11,463) areas. Conversely, small containers hosted higher proportions of immatures in the rural (38.34%, 4395/11463) than the suburban (25.62%, 3,745/14,617) and urban (17.00%, 3,902/22,949) areas. Medium containers provided over 10% of immatures in the suburban (18.93, 2767/14617) and rural (14.48%, 1,660/11,463) areas, but only 6.27% (1,438/22,949) in the urban areas. *Aedes aegypti* immatures were sampled among a large range of wet containers, but mostly in tires (46.16%, 22,632/49,029) and small containers (24.56%, 22,632/49,029) in all the study areas. In the urban areas, the most productive breeding sites of *Ae. aegypti* were tires (57.60%, 13,219/22,949), followed by small containers (15.04%, 3,451/22,949). However, *Ae. aegypti* was abundantly found within small containers in the suburban (17.30%, 2,301/13,300) and rural areas (20.15%, 2,574/12,780). Furthermore, large containers, medium containers, flower pots and others produced at least 1% of immatures in all the study areas. *Ades. lilii, Ae. luteocephalus* and *Tx. brevipalipi* were mainly collected among tires in the suburban areas and small containers in the rural areas. Tires were the most providers of *Cx. nebulosus* and *Cx. annulioris* larvae in all the study areas, with the highest productivity rates of 8.26% (901/14,617) and 6.16% (1,207/14,617) observed in the suburban areas, respectively.

#### Biological associations and interactions in containers

The present study revealed multiple biological associations among mosquito larvae dwelling in the same water-holding containers. These cohabitations and bio-interactions varied across the study areas and seasons. Fifteen positive and four negative associations were identified in all the study areas. The highest number of biological associations was recorded in the rural areas (nine associations), whereas the lowest number was observed in the urban areas (five associations). Among the positive associations, *Ae. aegypti* were frequently associated with *Cx. annulioris* and *Cx. nebulosus* immatures within tires and small containers in all the study areas. *Ae. aegypti* and *An. gambiae* larvae cohabited in the urban areas, during SDS. *Ae. aegypti, An. gambiae* and *Cx. nebulosus* co-existed in the same larval containers in the suburban areas during LDS. *Ae. aegypti* co-shared larval breeding sites with *Ae. dendrophilus* in LRS and *Ae. lilii* in SDS in rural areas. The negative associations observed involved predatory larvae of *Er. chrysogaster* with *Ae. aegypti*, *Lu. tigripes* with *Ae. aegypti, Cx. nebulosus,* and *Cx. annulioris* and *Ae. fraseri*, that mainly occurred in the rural areas. During SDS, *Lu. tigripes* larvae co-bred with *Ae. aegypti* and *An. gambiae* immatures in tires in the urban areas.

### Stegomyia indices

#### Container index

Overall, CI was estimated at 34.39% (95% CI: [33.07%, 35.70%]) in all the study areas (Fig 5). The highest CI was recorded in the urban areas (40.43%, 95% CI: [38.18%, 442.76%]), followed by the suburban (32.02%, 95% CI: [29.75%, 34.29%]) and rural (29.87%, 95% CI: [27.62, 32.12]) areas. CI values differed significantly among the study areas (F = 4.83, df = 2, p < 0.0001). CI values were significantly higher in the urban compared with the rural areas (F = 10.57, df = 2, p < 0.0001), but similar to the suburban areas (F = 9.11, df = 1, p = 0.056). CI values did not vary significantly in all the study areas across seasonal variations (F = 1.58, df = 11, p = 0.1698).

**Fig 5.**
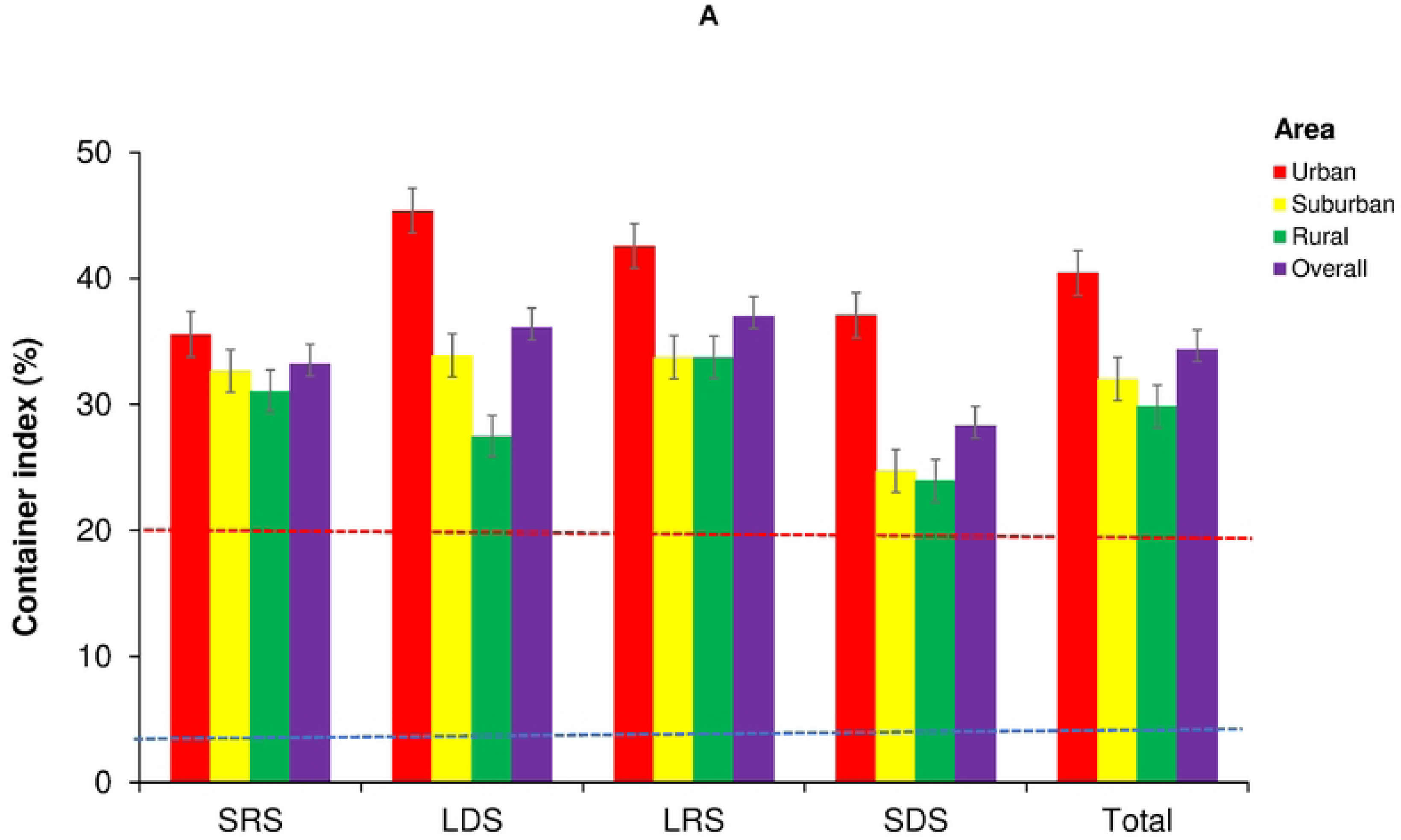
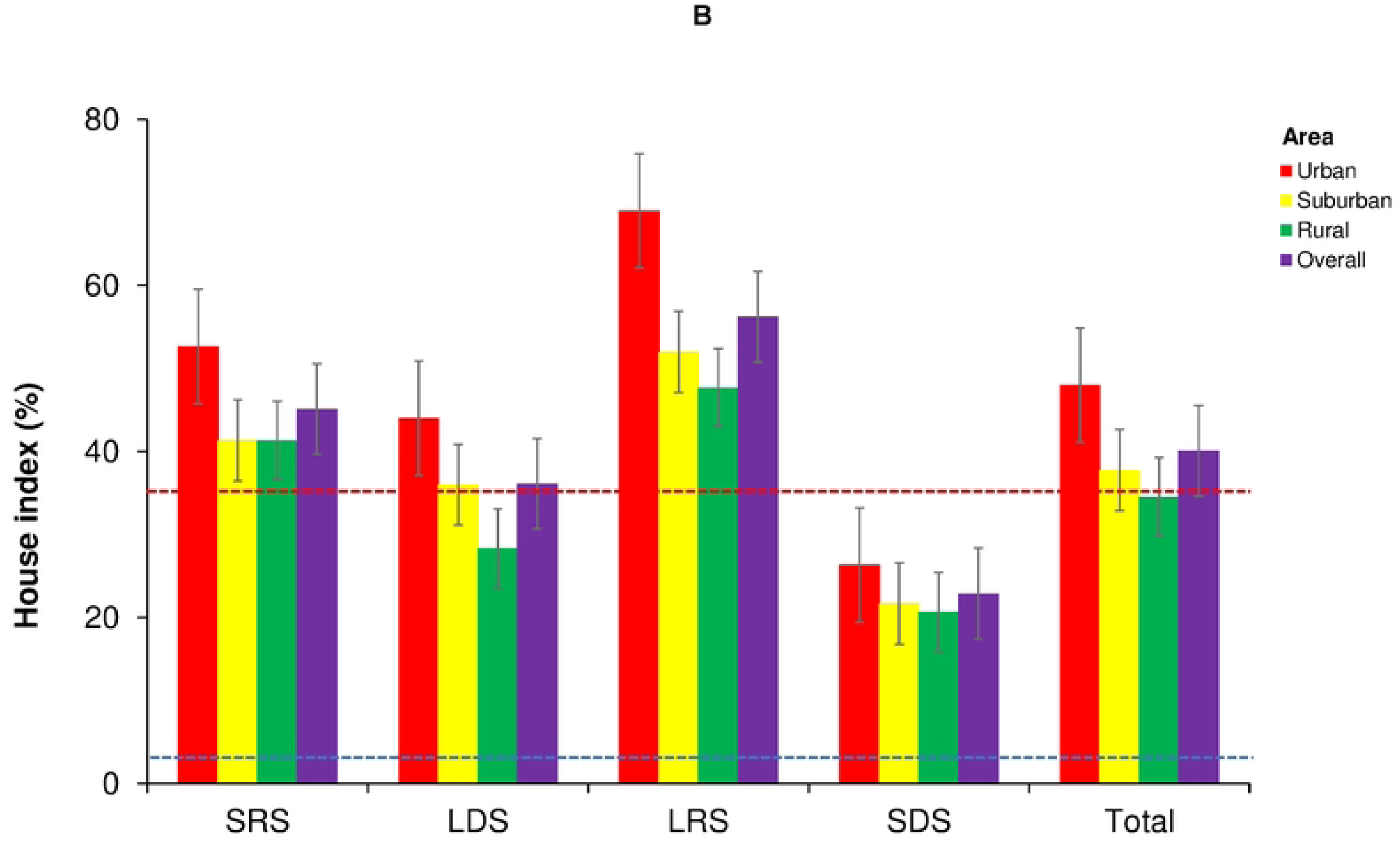
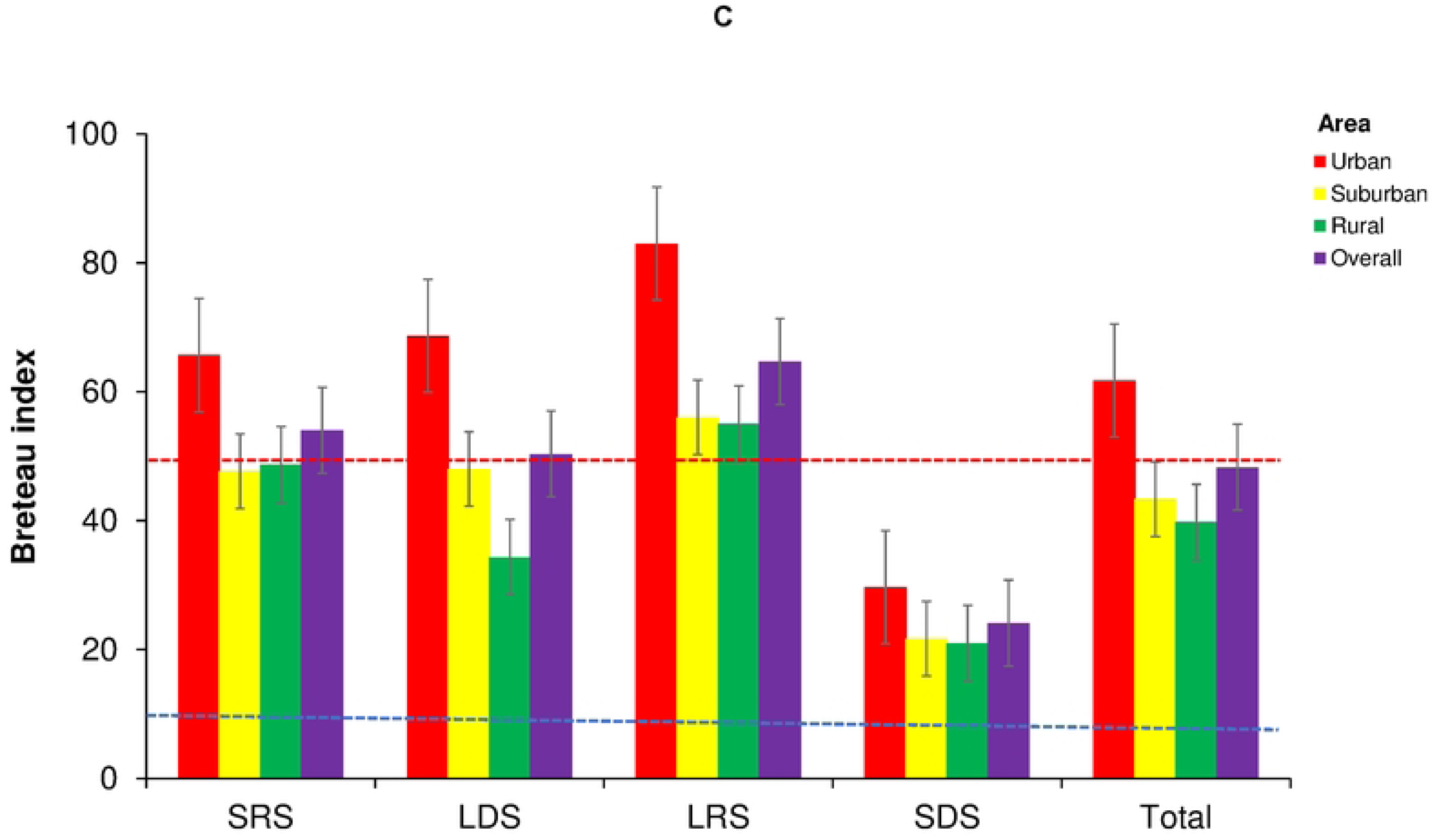
Seasonal variations in *Aedes aegypti* larval in across an urban-rural gradient in Cocody-Bingerville, southeastern Côte d’Ivoire from August 2023 to July 2024. Error bars show the confidence intervals (95% CI). A: House index (HI), B: Container index (CI), C: Breteau index (BI). SRS: short rainy season, LDS: long dry season, LRS: long rainy season, SDS: short dry season. The blue dotted lines represent the dengue epidemic thresholds. The red dotted lines represent the yellow fever epidemic thresholds. The dengue epidemic thresholds are 4% for house index, 3% for container index and 5 for Breteau index [50]. The yellow fever epidemic thresholds are 35% for house index, 20% for container index and 50 for Breteau index [51]. (PDF).

#### House index

HI was of 40.08% (95% CI: [34.32%, 45.85%]) in all the study areas (Fig 5). The urban areas had the highest HI values with 48.00% (95% CI: [35.22%, 60.78%]), followed by the suburban (37.75%, 95% CI: [27.71%, 47.79%]) and rural (34.50%, 95% CI: [26.41%, 42.59%]) areas. HI values were statistically comparable between all the study areas (F = 2.19, df = 2, p = 0.1279). In the urban areas, HI was higher during LRS (69.00%, 95% CI: [47.78%, 90.22%]) compared with SDS (26.33%, 95% CI: [-11.37%, 64.03%]) (F = −42.67, df = 1, p < 0.0001). In the suburban areas, HI value statistically was higher during LRS (52.00%, 95% CI: [34.61%, 69.39%]) compared with SDS (21.67%, 95% CI: [-10.65, 53.99]) (F = −41.00, df = 1, p < 0.0001). HI was significantly higher in the urban areas during LRS than that recorded during SDS (F = −54.33, df = 1, p < 0.0001) and LDS (36.00%, 95% CI: [-1.51, 73.51]) (F = 35.33, df = 1, p < 0.0001) in the suburban areas, and rural areas during SDS (21.00%, 95% CI: [-0.66, 42.66]) (F = −41.33, df = 1, p < 0.0001). Finally, the urban areas displayed statistically higher HI values during SRS (52.67%, 95% CI: [15.38, 89.96]) than in the suburban areas during SDS (F = 38.00, df = 1, p < 0.0001).

#### Breteau index

The overall value of BI was 48.28 (95% CI: [40.61, 55.94]) in all the study areas (Fig 5). The urban areas with showed the highest BI value of 61.75 (95% CI: [45.16, 78.34]) that was followed by 43.33 (95% CI: [30.60, 56.06]) in the suburban and 39.75 (95% CI: [29.77, 49.73]) in the rural areas. ANOVA showed that BI differed significantly among the study areas (F = 3.77, df = 2, p < 0.0001). BI values significantly different between the urban and rural areas only (F = 22.00, df = 1, p < 0.0001). BI values significantly varied across seasonality (F = 4.82, df = 11, p < 0.0001). In the urban areas, BI values were statistically higher during LRS (83.00, 95% CI: [69.85, 96.14]) compared with SDS in the suburban areas (21.67, 95% CI: [-10.66, 53.99]) (F = −68.00, df = 1, p < 0.0001) and the rural areas (21.00, 95% CI: [-0.66, 42.66]) (F = −55.33, df = 1, p < 0.0001). Moreover, BI in the urban areas recorded during LDS (68.67, 95% CI: [34.21, 103.12]) and SRS (65.67, 95% CI: [-8.04, 139.37]) were significantly higher than that found in the suburban areas during SDS (F = −53.67, df = 1, p < 0.0001 and F = 50.67, df = 1, p < 0.0001, respectively).

### Epidemic risk of dengue and yellow fever

The respective values of CI, HI and BI were 34.39%, 40.08%, and 48.28 in all the study areas, 40.43%, 48.00% and 61.75 in the urban, 32.02%, 37.75% and 43.33 in the suburban, and 29.87%, 34.50% and 39.75 in the rural areas (Table 6). According to the WHO density scale, these risk values corresponded to the intervals of 5–8 in the all study, 6–8 in the urban, 5–7 in the suburban and 5–7 in the rural areas. All risk indices exceeded the WHO DEN and YF epidemic thresholds for and in the areas [50,51]. Risk indices indicated high DEN risk in all the areas. YF risks were high in the urban and moderate in the suburban and rural areas. DEN epidemic risks were permanently high in all the study areas during all seasons, except for during SDS when the suburban areas showed medium risk. YF epidemic risks were moderate in all the study areas in almost all seasons, but high during LDS and LRS in the urban and LRS in the suburban areas.

**Table 6.** *Aedes aegypti* larval indices and risk of transmission of dengue and yellow fever viruses across an urban-rural gradient in the arboviral hotspots of Cocody-Bingerville, southeastern Côte d’Ivoire from August 2023 to July 2024. (PDF).

### Correlation between ovitrap indices and *Stegomyia* indices

All the correlations among ovitrap and *Stegomyia* indices were positive (S8 Table). For ovitrap indices, MEO was strongly and significantly correlated with OPI (ρ = 0.7709, p < 0.0001) and EDI (r = 0.7242, p < 0.0001). The correlation between OPI and EDI was weak and not significant (ρ = 0.1748, p = 0.3063). For Stegomyia indices, we detected very strong and significant correlations between all indices: CI and HI (r = 0.8467, p < 0.0001), CI and BI (r = 0.9413, p < 0.0001), and HI and BI (r = 0.9452, P < 0.0001). We found several relationships between ovitrap and *Stegomyia* indices. Indeed, OPI was moderately and significantly correlated with CI (ρ = 0.3967, p = 0.0172) and BI (ρ = 0.3582, p = 0.0325), and was weakly and not statistically correlated with HI (ρ = 0.2542, p = 0.1342). MEO showed moderate and significant correlations with HI (r = 0.3644, p = 0.0289) and BI (r = 0.3577, p = 0.0322), but was weakly and not statistically correlated with ci (r = 0.2856, p = 0.0913). EDI was weakly and not significantly correlated with CI (r = 0.1041, p = 0.5456), HI (r = 0.1936, p = 0.2580) and BI (r = 0.1749, p = 0.3077).

## Discussion

As most urbanized and urbanizing African countries, Côte d’Ivoire has recurrently faced multiple DEN outbreaks, often combined with YF cases recently, 2017-2024 [23,24, 28–31]. Over 80-90% DEN and YF cases of the country were reported in the health district of Cocody-Bingerville subjected to a rapid and uncontrolled urbanization [39,52]. To better understand the DEN and YF outbreak dynamics in the region we assessed the oviposition and larval ecologies of mosquitoes and ovitrap indices and *Aedes Stegomyia* indices along an urban-rural gradient (i.e., in the urban, suburban and rural areas) during ongoing outbreaks within the arboviral hotspots in Cocody-Bingerville. The results demonstrated that all the study areas were highly infested with diversified mosquito species, including wild *Aedes* vector, sympatric and predatory species mostly confined to rural and suburban areas, and widely dominated by the main arbovirus vector *Ae. aegypti* that was primarily abundant (>90% of mosquito fauna) in the urban areas. *Stegomyia* indices were above the WHO DEN and YF epidemic thresholds in all the areas, and showed that people were much more exposed to DEN outbreaks in the urban areas. These results are important for understanding and developing more sustainable programs to control the current DEN outbreaks.

Our data showed that all the study areas had high capacity for hosting large numbers and high diversity mosquito immatures (70,550 specimens and 15 species). Importantly, confirmed or potential mosquito vectors of arboviruses were commonly found in all the study areas. Indeed, our collections yielded several *Aedes* and non-*Aedes* mosquito species that can host or transmit DENV, YFV and other viruses of the public and global health relevance [16]. Collected *Aedes* mosquitoes included *Ae. aegypti*, *Ae. luteocephalus*, *Ae. metallicus*, and *Ae. vittatus* have been shown to carry and/or to transmit in nature over 24 viruses, including DENV, YFV), CHIKV, ZIKV, and RVFV in tropical regions [10,11]. As *Ae. dendrophilus*, *Ae. fraseri, Ae. lilii* belong to the same *Stegomyia* subgenus as *Ae. aegypti* the main vector of DENV and YFV, they could be suspected as potential vectors of DENV and YFV, as well. Among non-*Aedes* mosquitoes, *Er. Chrysogaster* have been found to have natural infection, while *Er. quinquevittatus* has exhibited laboratory competence with YFV in Africa [9]. CHIKV and O’nyong-nyong virus has been isolated from the malaria *An. gambiae* vector [53]. *Culex quinquefasciatus* has been shown susceptible to transmit RVFV [54]. The presence of numerous vectors, including anthropophagic and non-anthropophagic species, in the suburban and rural areas suggest the probable co-existence of multiple cycles and co-circulations of still unidentified arboviruses, in addition to DENV and YFV in the ecosystems of Cocody-Bingerville, as suggested by Weaver et al. [55]. Wild *Aedes* vectors present in the domestic ecozones may be involved into the epizootic cycles acting as bridge vectors of DENV and YFV at human-wildlife interface, with possible spillovers (wild animal-to-human) and spillbacks (human-to-wild/domestic animal), while the sylvatic vectors present in the peridomestic ecozones near forests or farms might be involved in the enzootic cycles (wild animal-to-wild animals) in the suburban and rural areas [56,57]. The highly anthropophagic *Ae. aegypti* vector could amply human-to-human epidemic transmission of DENV and YFV in the urban areas [55].

Our data demonstrated that mosquito larval breeding sites and biological interactions shaped across the urbanization levels. We found mosquito immatures in both natural (e.g., banana leaf axils, tree holes, bamboo holes and snail shells) and artificial containers (e.g., flowerpots, tires, abandoned containers and water storage tanks) habitats in both domestic and peridomestic ecozones in the rural and suburban areas. We detected in these water-holding containers several biological associations among *Aedes* and non-*Aedes* larvae that could act as competitors (*Aedes* and non-*Aedes* immatures) and predators (*Eretmapodites*, *Lutzia tigripes* and *Toxorhynchites*) that appear to regulate *Aedes* species diversity, and reduce *Ae. aegypti* density and DENV and YFV epidemic risks in the suburban and rural areas, as previously reported in rural Côte d’Ivoire [16] and Senegal [58, 59]. The predatory larvae could serve as biological agents for biocontrol of *Aedes* vectors for arboviral prevention [19]. The larvae of sympatric *Culex* species, such as *Cx. quinquefasciatus*, were associated with *Ae. aegypti* presence in containers in urban Côte d’Ivoire [19]. Additionally, *An. gambiae* was unexpectedly sampled with *Aedes* in artificial containers in the urban areas. The presence of *An. gambiae* larvae in urban containers may derive from new ovipositional adaption to survive from absence or rarity of their usual breeding sites (e.g., puddles) [60,61], and should require a particular attention in the current context of rapid spread and invasion of similar container-specialist, *Anopheles stephensi,* a competent malaria vector across Africa [62,63]*. Anopheles stephensi* has been reported in Ghana, close to Cocody-Bingerville [62,63].

Our results indicated that *Ae. aegypti* was the most prevalent species in all the study areas, spread from rural to urban areas, and showed the highest densities in the urban areas. *Ae. aegypti* populations may have probably taken advantage of the reduction of predatory and competitive pressure and agricultural chemical pesticides coupled with an increase in the numbers of man-made containers and human hosts across increasing urbanization to proliferate in the urban areas and the domestic ecozones. The increased numbers of the artificial breeding containers (e.g., tires, small containers, etc.) and human hosts in the urban areas and domestic ecozones offer greater ovipositing and blood-feeding opportunities to *Ae. aegypti* females. Previous studies have shown that the urban areas of Cocody-Bingerville harbor large proportions of *Ae. aegypti* populations and larval breeding sites (discarded containers, tires, water storage containers, etc.) due to a poor environmental sanitation and hygiene [16]. Additionally, *Ae. aegypti* propositions were higher in the domestic compared with peri-domestic ecozones in the suburban and rural areas, but similar between the two ecozones in the urban areas. The high presence of *Ae. aegypti* within and around houses may be explained by their strong anthropophilic behaviors that could increase the risks of DENV and YFV transmission to the local residents [16,20,36,64,65].

Our results revealed that *Ae. aegypti* oviposition indices and *Stegomyia* indices gradually increased along increasing urbanization gradient, and reached the highest values in the urban areas thus incriminating them as the riskiest areas. Indeed, the significantly highest values of ovitrap indices (OPI, MEO and EDI) and *Stegomyia* indices (CI, HI and BI) were found in the urban areas. All *Stegomyia* indices exceeded the WHO DEN and YF epidemic thresholds in all study areas, and corresponded to the WHO risk intervals of 5–8, 6–8, 5–7 and 5–7 in all the study, urban, suburban and rural areas, respectively. The levels of risk indices indicate that DEN risk was high in all the study areas, whereas YF risk was high in the urban and moderate in suburban and rural areas. However, the massive presence of numerous sylvatic *Aedes* species and competent vectors (*Ae. vittatus* and *Ae. luteocephalus*) in the suburban and rural areas, in addition to *Ae. aegypti*, is expected to increase DENV and YFV transmission risks in these non-urban settings. The risk levels of DEN and YF were less influenced by seasonality [36,40], but mostly by urbanization. DEN epidemic risks were permanently high in all the study areas and all seasons, and medium only during SDS in the suburban areas. Moreover, YF epidemic risks were much more moderate in all the study areas across the seasons, and found high only during LDS and LRS in the urban and LRS in the suburban areas [36,40]. Above all, *Stegomyia* indices were all largely and permanently above the WHO DEN epidemic thresholds in all areas and during all seasons, and this could explain the recurrent and ongoing DEN outbreaks in Cocody Bingerville. Our study showed that the ovitrap indices (OPI, MEO and EDI) and *Stegomyia* indices (CI, HI and BI) were almost correlated, suggesting that the oviposition indices were predictive of larval indices. Additionally, OPI, MEO and EDI were correlated each with other, as well as CI, HI and BI. Both egg and larval data produced valuable information on the spatio-temporal distribution of *Ae*. *aegypti* and DEN and YF risks at different urbanization levels. The current study used, for the first time in Côte d’Ivoire, OPI, MEO and EDI indicators to assess *Aedes* oviposition activities. The correlations between ovitrap indices and *Stegomyia* indices suggest that ovitrap indicators could be appropriated for the assessment of DEN and YF epidemic risks in Côte d’Ivoire. The use of ovitrap indices could offer new opportunities and complementary resources for surveillance and control of *Aedes* DEN and YF vectors and risks. Indeed, ovitrap-based surveillance is simpler in use, less resource-demanding, less time-consuming and cheaper method compared with larval survey method. Thus, ovitrap might provide valuable resources for public health and research in arboviral surveillance, suffering from lack of financial and scientific investments in Africa [3,4].

While *Stegomyia* indices contributed to better understanding the entomological mechanisms of the ongoing DEN outbreaks in urban Cocody-Bingerville, they were not fully predictive of DEN and YF outbreaks in the suburban and rural areas. This lack of correlations between *Stegomyia* indices and DEN and YF outbreaks could be probably due to some methodological and logistically limitations and biases that need be addressed. Indeed, even risk levels were moderate in the suburban and rural areas, they all were above the WHO epidemic thresholds for DEN and YF in these areas, but no outbreaks were reported there during the study period, 2023-2024. The DEN and YF outbreaks were mainly reported in urban areas from 2023 to 2024 and in previous years. This suggests that the high abundances of *Ae. aegypti* and other *Aedes* species and high epidemic risk indices only could not be sufficient to trigger an outbreak of DEN or YF, as observed in urban areas of Côte d’Ivoire [36], Burkina Faso [66] and Kenya [67]. Moreover, DEN infections are mostly misdiagnosed as malaria or recorded as non-malaria febrile illnesses or indetermined fevers [68,69], biasing the disease prevalence as the incidences are mainly captured as clinical cases in hospitals [68]. Complementary investigations focusing on the detection of DEN and YF viruses and/or anti-bodies in the whole human populations and in *Aedes* vectors in the present study areas could help to better understand the epidemiological dynamics DEN and YF there. The presence of the sylvan *Aedes* species could underestimate arboviral risk measured by *Stegomyia* indices, and should be taken into account when interpreting the risk levels, mainly in rural areas. Further studies on the host-seeking, blood-feeding and resting behaviors and related genetic mechanisms of local *Aedes aegypti* adults in domestic *versus* peridomestic ecozones, indoors *versus* outdoors, and water storage *versus* discarded containers could be recommended. Investigating the socio-cultural and behavioral factors in the local communities relative to the management of waste and water containers, and awareness about *Aedes* and DEN and YF risks would provide added value for developing sustainable community-based vector control programs for effective prevention these diseases in the health district of Cocody-Bingerville.

In summary, our findings revealed that mosquito species composition and abundance shift at different levels of urbanization towards the spread and the dominance of *Ae. aegypti* in the urban areas. As in many other African regions, urbanization accelerates at a fast and uncontrolled pace in the health district of Cocody-Bingerville, advantaging the urban *Ae. aegypti* species when restricting wild *Aedes* species to the suburban and rural areas. This tendency is paralleled to increased risks and multiple resurgences of DEN and YF, and may thus explain the current DEN outbreaks in urban Cocody-Bingerville, while suburban and rural areas are also exposed to high epidemic risks. Therefore, while strengthened warning systems and operated vector control interventions (e.g., community awareness and mobilization, removals of larval breeding sites and insecticide space sprays against *Aedes* [28]), are understandably performed in the urban areas of Cocody-Bingerville as usual, the suburban and rural areas should be considered when developing and applying vector programs as these areas could act as reservoirs and (re-)emergence foci due to the presence of several sylvatic bridge vectors susceptible of spillover and spillback of DENV and YFV in the study region [56,57]. Our study provides a baseline for raising local community awareness and guiding appropriate preventative actions, such as community-based management of the identified key *Aedes* larval sites (i.e., tires and small containers). Suggested *Aedes* vector actions should be inclusive and holistic interventions planned and conducted in a multisectoral framework [70–73]. Such an integrated program should involve public engagement and mobilization of government and local policy-makers, urban planners from public and private sectors, local communities and scientists for joint efforts for a sustainable reduction of key larval habitats of *Aedes* vectors and arboviral risks to constraint ongoing DEN outbreaks and prevent future arboviral epidemics in the health district of Cocody-Bingerville southern Côte d’Ivoire.

## Conclusions

Our results improve our understanding of how urbanization shapes the species biodiversity and larval breeding sites of mosquito immatures and the risks of transmission of DEN and YF within arboviral hotspots in the health district of Cocody-Bingerville, southeastern Côte d’Ivoire. Our findings demonstrated that increasing urbanization was associated with a regression of species richness and a restriction of wild *Aedes* species in the rural and suburban areas, and a strong dominance of *Ae. aegypti* and raised DEN and YF risks in the urban areas. Wild *Aedes* species could act as bridge vectors involved into spillover and spillback cycles in the rural and suburban areas, while *Ae. aegypti* might transmit DENV and YFV to people leading to epidemics in the urban areas. Larvae mostly bred in tires in the urban and suburban and small containers in the rural areas. Ovitrap indices corelated with *Stegomyia* indices that were above the WHO DEN and YF epidemic thresholds in all the areas, the highest risk levels recorded in the urban areas. The results suggest that the people are exposed to high DEN risk and moderate YF risk in all the areas. This may explain the recurrent and ongoing DEN outbreaks in the urban settings of Cocody-Bingerville. Our study provides a baseline for raising local community awareness and guiding appropriate preventative actions, including a community-based management of identified key larval habitats to control ongoing DEN outbreaks in Côte d’Ivoire and in other similar African settings

## Acknowledgements

The authors thank the administrative, health and traditional authorities who authorized the research activities, the mosquito and data collectors, and the inhabitants of all the study areas.

## Author contributions

**Conceptualization:** JZBZ, GDM, LSD, JFM, AAA, SB, SCB.

**Data curation:** YNB, JZBZ, JDKD, PNC.

**Formal analysis:** YNB, JZBZ.

**Funding acquisition:** JZBZ, GDM, LSD, JFM, AAA, SB, SCB.

**Investigation:** YNB, JZBZ, JDKD, PNC, MT.

**Methodology:** YNB, JZBZ, PNC, PGM, FH, LSD, JFM, SCB, MT.

**Project administration:** JZBZ, FH, SB, SCB.

**Data collection:** YNB, JDKD, PNC.

**Resources:** JZBZ, MT.

**Supervision:** JZBZ, MT.

**Visualization:** YNB, JZBZ, JDKD.

**Writing – original draft:** YNB, JZBZ.

**Writing – review & Editing:** YNB, JZBZ, PNC, PGM, FH, GDM, LSD, JFM, AAA, SB, SCB, MT.

## Abbreviations

BI: Breteau Index
CI: Container Index
DEN: dengue
DENV: dengue virus
EDI: egg density index
HI: House Index
LDS: long dry season
LRS: long rainy season
MEO: mean egg counts per ovitrap
OPI: ovitrap positivity index
SDS: short dry season
SRS: short rainy season
WHO: World Health Organization
YF: yellow fever
YFV: yellow fever virus.

## Supporting information captions

**S1 Table. Geographical distribution of adult mosquitoes emerged from eggs and larvae sampled among domestic and peridomestic ecozones along an urban-rural gradient in the arboviral hotspots of Cocody-Bingerville, southeastern Côte d’Ivoire from August 2023 to July 2024. (PDF).**

**S2 Table. *Aedes aegypti* oviposition indices across an urban-rural gradient in Cocody-Bingerville, southeastern Côte d’Ivoire from August 203 to July 2024. (PDF).**

**S3 Table. Outputs of data analysis comparing the ovitrap positivity index of *Aedes aegypti* along an urban-rural gradient in Cocody-Bingerville, southeastern Côte d’Ivoire from August 203 to July 2024. Results are the outputs of the generalized linear mixed model (GLMM) procedures. Results are considered significant for p-values < 0.05. (PDF).**

**S4 Table. Outputs of data analysis comparing the mean egg count per ovitrap of *Aedes aegypti* along an urban-rural gradient in Cocody-Bingerville, southeastern Côte d’Ivoire from August 203 to July 2024. Results are the outputs of the generalized linear mixed model (GLMM) procedures. Results are considered significant for p-values < 0.05. (PDF).**

**S5 Table. Outputs of data analysis comparing the egg density index of *Aedes aegypti* along an urban-rural gradient in Cocody-Bingerville, southeastern Côte d’Ivoire from August 203 to July 2024. Results are the outputs of the generalized linear mixed model (GLMM) procedures. Results are considered significant for p-values < 0.05. (PDF).**

**S6 Table. Seasonal variations in the positivity of the breeding sites of mosquitoes across an urban-rural gradient in the arboviral hotspots of Cocody-Bingerville, southeastern Côte d’Ivoire from August 2023 to July 2024. (PDF).**

**S7 Table. Distribution of mosquito species according to categories of breeding sites in along an urban-rural gradient in the arboviral hotspots of Cocody-Bingerville, southeastern Côte d’Ivoire from August 2023 to July 2024. (PDF).**

**S8 Table. Outputs of data analysis of relationship between oviposition indices and *Stegomyia* indices of *Aedes aegypti* along an urban-rural gradient in Cocody-Bingerville, southeastern Côte d’Ivoire from August 203 to July 2024. (PDF).**

## References

1. Gubler DJ. Dengue and dengue hemorrhagic fever. Clin Microbiol Rev. 1998 Jul;11(3):480–96. doi: 10.1128/CMR.11.3.480. PMID: 9665979; PMCID: PMC88892.

2. LaBeaud AD, Sutherland LJ, Muiruri S, Muchiri EM, Gray LR, Zimmerman PA, Hise AG, King CH. Arbovirus prevalence in mosquitoes, Kenya. Emerg Infect Dis. 2011 Feb;17(2):233–41. doi: 10.3201/eid1702.091666. PMID: 21291594; PMCID: PMC3204744.

3. Weetman D, Kamgang B, Badolo A, Moyes CL, Shearer FM, Coulibaly M, Pinto J, Lambrechts L, McCall PJ. Aedes Mosquitoes and Aedes-Borne Arboviruses in Africa: Current and Future Threats. Int J Environ Res Public Health. 2018 Jan 28;15(2):220. doi: 10.3390/ijerph15020220. PMID: 29382107; PMCID: PMC5858289.

4. Buchwald AG, Hayden MH, Dadzie SK, Paull SH, Carlton EJ. Aedes-borne disease outbreaks in West Africa: A call for enhanced surveillance. Acta Trop. 2020 Sep;209:105468. doi: 10.1016/j.actatropica.2020.105468. Epub 2020 May 19. PMID: 32416077.

5. Mordecai EA, Ryan SJ, Caldwell JM, Shah MM, LaBeaud AD. Climate change could shift disease burden from malaria to arboviruses in Africa. Lancet Planet Health. 2020 Sep;4(9):e416–e423. doi: 10.1016/S2542-5196(20)30178-9. PMID: 32918887; PMCID: PMC7490804.

6. Agboli E, Tomazatos A, Maiga-Ascofaré O, May J, Lühken R, Schmidt-Chanasit J, Jöst H. Arbovirus Epidemiology: The Mystery of Unnoticed Epidemics in Ghana, West Africa. Microorganisms. 2022 Sep 27;10(10):1914. doi: 10.3390/microorganisms10101914. PMID: 36296190; PMCID: PMC9610185.

7. Africa Urban Forum: co-creating solutions to make cities habitable for the growing population https://au.int/en/pressreleases/20240904/africa-urban-forum-co-creating-solutions-make-cities-habitable-growing. Accessed 20 December 2024.

8. Qi X, Wang Y, Li Y, Meng Y, Chen Q, Ma J, Gao GF. The Effects of Socioeconomic and Environmental Factors on the Incidence of Dengue Fever in the Pearl River Delta, China, 2013. PLoS Negl Trop Dis. 2015 Oct 27;9(10):e0004159. doi: 10.1371/journal.pntd.0004159. PMID: 26506616; PMCID: PMC4624777.

9. Braack L, Gouveia de Almeida AP, Cornel AJ, Swanepoel R, de Jager C. Mosquito-borne arboviruses of African origin: review of key viruses and vectors. Parasit Vectors. 2018 Jan 9;11(1):29. doi: 10.1186/s13071-017-2559-9. PMID: 29316963; PMCID: PMC5759361.

10. Rose NH, Sylla M, Badolo A, Lutomiah J, Ayala D, Aribodor OB, Ibe N, Akorli J, Otoo S, Mutebi JP, Kriete AL, Ewing EG, Sang R, Gloria-Soria A, Powell JR, Baker RE, White BJ, Crawford JE, McBride CS. Climate and Urbanization Drive Mosquito Preference for Humans. Curr Biol. 2020 Sep 21;30(18):3570–3579.e6. doi: 10.1016/j.cub.2020.06.092. Epub 2020 Jul 23. PMID: 32707056; PMCID: PMC7511451.

11. Kolimenakis A, Heinz S, Wilson ML, Winkler V, Yakob L, Michaelakis A, Papachristos D, Richardson C, Horstick O. The role of urbanisation in the spread of Aedes mosquitoes and the diseases they transmit-A systematic review. PLoS Negl Trop Dis. 2021 Sep 9;15(9):e0009631. doi: 10.1371/journal.pntd.0009631. PMID: 34499653; PMCID: PMC8428665.

12. Wilke ABB, Vasquez C, Carvajal A, Moreno M, Fuller DO, Cardenas G, Petrie WD, Beier JC. Urbanization favors the proliferation of Aedes aegypti and Culex quinquefasciatus in urban areas of Miami-Dade County, Florida. Sci Rep. 2021 Nov 26;11(1):22989. doi: 10.1038/s41598-021-02061-0. PMID: 34836970; PMCID: PMC8626430.

13. Agha SB, Tchouassi DP. Urbanization of Aedes mosquito populations and evolution of arboviral disease risk in Africa. Curr Opin Insect Sci. 2022 Dec;54:100988. doi: 10.1016/j.cois.2022.100988. Epub 2022 Nov 1. PMID: 36332839.

14. Poongavanan J, Lourenço J, Tsui JL, Colizza V, Ramphal Y, Baxter C, Kraemer MUG, Dunaiski M, de Oliveira T, Tegally H. Dengue virus importation risks in Africa: a modelling study. Lancet Planet Health. 2024 Dec;8(12):e1043–e1054. doi: 10.1016/S2542-5196(24)00272-9. PMID: 39674194; PMCID: PMC11649930.

15. World Health Organization (21 December 2023). Disease Outbreak News; Dengue – Global situation Available at: https://www.who.int/emergencies/disease-outbreak-news/item/2023-DON498. Accessed 22 December 2024.

16. Zahouli JBZ, Koudou BG, Müller P, Malone D, Tano Y, Utzinger J. Urbanization is a main driver for the larval ecology of Aedes mosquitoes in arbovirus-endemic settings in south-eastern Côte d’Ivoire. PLoS Negl Trop Dis. 2017 Jul 13;11(7):e0005751. doi: 10.1371/journal.pntd.0005751. PMID: 28704434; PMCID: PMC5526600.

17. Yang B, Borgert BA, Alto BW, Boohene CK, Brew J, Deutsch K, DeValerio JT, Dinglasan RR, Dixon D, Faella JM, Fisher-Grainger SL, Glass GE, Hayes R Jr, Hoel DF, Horton A, Janusauskaite A, Kellner B, Kraemer MUG, Lucas KJ, Medina J, Morreale R, Petrie W, Reiner RC Jr, Riles MT, Salje H, Smith DL, Smith JP, Solis A, Stuck J, Vasquez C, Williams KF, Xue RD, Cummings DAT. Modelling distributions of Aedes aegypti and Aedes albopictus using climate, host density and interspecies competition. PLoS Negl Trop Dis. 2021 Mar 25;15(3):e0009063. doi: 10.1371/journal.pntd.0009063. PMID: 33764975; PMCID: PMC8051819.

18. Lizuain AA, Maffey L, Garzón M, Leporace M, Soto D, Diaz P, Salomón OD, Santini MS, Schweigmann N. Larval Competition Between Aedes albopictus and Aedes aegypti (Diptera: Culicidae) in Argentina: Coexistence and Implications in the Distribution of the Asian Tiger Mosquito. J Med Entomol. 2022 Sep 14;59(5):1636–1645. doi: 10.1093/jme/tjac102. Erratum in: J Med Entomol. 2023 Sep 12;60(5):1136-1137. doi: 10.1093/jme/tjad080. PMID: 35899788.

19. Sukupayo PR, Poudel RC, Ghimire TR. Nature’s Solution to *Aedes* Vectors: *Toxorhynchites* as a Biocontrol Agent. J Trop Med. 2024 Jun 21;2024:3529261. doi: 10.1155/2024/3529261. PMID: 38948015; PMCID: PMC11213640.

20. Zahouli JB, Utzinger J, Adja MA, Müller P, Malone D, Tano Y, Koudou BG. Oviposition ecology and species composition of Aedes spp. and Aedes aegypti dynamics in variously urbanized settings in arbovirus foci in southeastern Côte d’Ivoire. Parasit Vectors. 2016 Sep 29;9(1):523. doi: 10.1186/s13071-016-1778-9. PMID: 27682270; PMCID: PMC5041276.

21. Zahouli JBZ, Koudou BG, Müller P, Malone D, Tano Y, Utzinger J. Effect of land-use changes on the abundance, distribution, and host-seeking behavior of Aedes arbovirus vectors in oil palm-dominated landscapes, southeastern Côte d’Ivoire. PLoS One. 2017 Dec 7;12(12):e0189082. doi: 10.1371/journal.pone.0189082. PMID: 29216248; PMCID: PMC5720743.

22. Chathuranga WGD, Weeraratne TC, Abeysundara SP, Karunaratne SHPP, de Silva WAPP. Breeding site selection and co-existing patterns of tropical mosquitoes. Med Vet Entomol. 2023 Sep;37(3):550–561. doi: 10.1111/mve.12656. Epub 2023 Apr 15. PMID: 37060294.

23. World Health Organization (WHO). Weekly Bulletin on Outbreaks and Other Emergencies— Week 42 2019, 14–20 October 2019. Geneva, Switzerland; 2019. Accessed 22 December 2024.

24. Côte d’Ivoire: Une épidémie de fièvre jaune touche 89 personnes, une personne décédée. Available: https://www.20minutes.fr/monde/2575051-20190731-cote-ivoire-epidemie-fievre-jaune-touche-89-personnes-personne-decedee. Accessed 22 December 2024.

25. Recensement Général de la Population et de l’Habitat (RGPH 2021). https://plan.gouv.ci/assets/fichier/RGPH2021-RESULTATS-GLOBAUX-VF.pdf. Accessed 28 December 2024.

26. ONU-Habitat Côte d’Ivoire Rapport pays | 2023. https://unhabitat.org/sites/default/files/2023/07/presentation_programme_cote_d_ivoire_fr.pdf. Accessed 28 December 2024.

27. Kone AB, Konan YL, Coulibaly ZI, Fofana D, Guindo-Coulibaly N, Diallo M, Doannio JM, Ekra KD, Odehouri-Koudou P. Évaluation entomologique du risque d’épidémie urbaine de fièvre jaune survenue en 2008 dans le district d’Abidjan, Côte d’Ivoire [Entomological evaluation of the risk of urban outbreak of yellow fever in 2008 in Abidjan, Côte d’Ivoire]. Med Sante Trop. 2013 Jan-Mar;23(1):66–71. French. doi: 10.1684/mst.2013.0153. PMID: 23693032.

28. Dengue fever – Côte d’Ivoire. Available: https://www.who.int/emergencies/disease-outbreak-news/item/04-august-2017-dengue-cote-d-ivoire-en. Accessed 22 December 2024

29. Côte d’Ivoire - Dengue outbreak 2023. In: FluTrackers News and Information. Available: https://flutrackers.com/forum/forum/emerging-diseases-other-healththreats-alphabetical-a-thru-h/dengue/976705-co^te-d-ivoire-dengue-outbreak-2023. Accessed 22 December 2024

30. CICG. Santé : tout savoir sur la lutte contre la dengue, ce mardi 23 juillet 2024. In: GOUV.CI [Internet]. Available: http://www.gouv.ci/_actualite-article.php?recordID=17219. Accessed 22 December 2024

31. Actualités Mes Vaccins. Available: https://www.mesvaccins.net/web/news/22376-la-dengue-en-afrique-bilan-des-signalements-faits-en-septembre-2024. Accessed 22 December 2024

32. Sylla Y, Diane MK, Adjogoua VE, Kadjo H, Dosso M. Dengue Outbreaks in Abidjan: Seroprevalence and Cocirculating of Three Serotypes in 2017. OSIR. 2021; 14. 10.59096/osir.v14i3.262527

33. Carlson R. Urban Yellow Fever Outbreak Infects 89 Causing One Fatality. 2019. Available: https://www.vaxbeforetravel.com/republic-c^ote-divoire-ivory-coast-reports-yellow-fevervirus-outbreak-abidjan. Accessed 22 December 2024.

34. Tropicale AS. Épidémie de Dengue : L’INHP détruit les gîtes larvaires de Cocody. Available: https://www.santetropicale.com/sites_pays/actus.asp?id=32045&action=lire&rep=rci. Accessed 22 December 2024.

35. WHO African Region Health Emergency Situation Report - Multi-country Outbreak of Dengue, Consolidated Regional Situation Report # 004 - Weekly bulletin on outbreaks and other emergencies, week 29: 15 to 21 July 2024 data as reported by: 17:00; 21 July 2024. https://reliefweb.int/report/ethiopia/weekly-bulletin-outbreaks-and-other-emergencies-week-29-15-21-july-2024-data-reported-1700-21-july-2024. Assessed 22 December 2024.

36. Adjobi CN, Zahouli JZB, Guindo-Coulibaly N, Ouattara AF, Vavassori L, Adja MA. Assessing the ecological patterns of Aedes aegypti in areas with high arboviral risks in the large city of Abidjan, Côte d’Ivoire. PLoS Negl Trop Dis. 2024 Nov 18;18(11):e0012647. doi: 10.1371/journal.pntd.0012647. PMID: 39556613; PMCID: PMC11611265.

37. Pan American Health Organization. Technical document for the implementation of interventions based on generic operational scenarios for Aedes aegypti control. Washington, D.C.: PAHO; 2019. https://iris.paho.org/handle/10665.2/51652. Assessed 22 December 2024.

38. WHO. Entomological surveillance for Aedes spp. in the context of Zika virus: interim guidance for entomologists. https://iris.who.int/bitstream/handle/10665/204624/WHO_ZIKV_VC_16.2_eng.pdf. Assessed 22 December 2024.

39. Fofana D, Beugré JMV, Yao-Acapovi GL, Lendzele SS. Risk of Dengue Transmission in Cocody (Abidjan, Ivory Coast). J Parasitol Res. 2019 Jan 14;2019:4914137. doi: 10.1155/2019/4914137. PMID: 30755798; PMCID: PMC6348904.

40. Guindo-Coulibaly N, Kpan MDS, Adja AM, Kouadio AMN, Assouho KF, Zoh DD, Azongnibo KRM, Remoue F, Fournet F. Seasonal variation and intra urban heterogeneity of the entomological risk of transmission of dengue and yellow fever in Abidjan, Côte d’Ivoire. Med Vet Entomol. 2022 Sep;36(3):329–337. doi: 10.1111/mve.12571. Epub 2022 Mar 30. PMID: 35352845.

41. Guindo-Coulibaly N, Adja AM, Coulibaly JT, Kpan MDS, Adou KA, Zoh DD. Expansion of Aedes africanus (Diptera: Culicidae), a sylvatic vector of arboviruses, into an urban environment of Abidjan, Côte d’Ivoire. J Vector Ecol. 2019 Dec;44(2):248–255. doi: 10.1111/jvec.12356. PMID: 31729805.

42. Cordellier R, Germain M, Hervy JP, Mouchet J. Guide pratique pour l’étude des vecteurs de fièvre jaune en Afrique et méthodes de lutte. 33rd ed. Paris: ORSTOM; 1977. http://www.documentation.ird.fr/hor/fdi:08619. Accessed 22 December 2024.

43. Cornet JP, Kittayapong P, Gonzalez JP. Le risque de transmission d’arbovirus par les tiques en Thailande [Risk of arbovirus transmission by ticks in Thailand]. Med Trop (Mars). 2004;64(1):43–9. French. PMID: 15224557.

44. Manrique-Saide P, Che-Mendoza, AzaelRizzo N, Pilger D, Lenhart A, Kroeger A, Arana B. Operational guide for assessing the productivity of Aedes aegypti breeding sites. World Heal Organ. 2011.

45. Rueda LM. Pictorial keys for the identification of mosquitoes (Diptera: Culicidae) associated with Dengue Virus Transmission. Zootaxa. 2004;589:1. doi:10.11646/zootaxa.589.1.1

46. van Strien AJ, Soldaat LL, Greory RD. Desirable mathematical properties of indicators for biodiversity change. Ecol Indic. 2012; 14: 202–208.

47. Protectus US. Student handout 1A: how to calculate biodiversity. http://entnemdept.ifas.ufl.edu/hodges/ProtectUs/lp_webfolder/9_12_grade/Student_Handout_1A.pdf; Assessed 22 December 2024.

48. Gomes AC. Medidas Dos Níveis de Infestação Urbana Para Aedes (Stegomyia) Aegypti Aedes (Stegomyia) Albopictus Em Programa de Vigilancia Entomológica. Inf. Epidemiológico SUS Braz. 1998, 7, 49–57.

49. Repeated measures analysis with Stata. http://www.ats.ucla.edu/stat/stata/seminars/repeated_measures/repeated_measures_analysis_stata.htm. Accessed 08 October 2024.

50. Pan American Health Organisation. Dengue and dengue hemorrhagic fever in the Americas: guidelines for prevention and control. Washington DC; 1994

51. World Health Organization (1971). Technical guide for a system of yellow fever surveillance = guide technique pour l’établissement d’un système surveillance de la fièvre jaune. Weekly Epidemiological Record = Relevé épidémiologique hebdomadaire, 46 (49), 493–500. https://iris.who.int/handle/10665/ 218521

52. Ano AKMN, Coulibaly D, Bénie VJ, Akani BC, Douba A, Ahoussou EM, et al. 411 – Situation épidémiologique de la dengue en Côte d’Ivoire de 2017 à 2020. Revue d’Epidémiologie et de Santé Publique. 2022; 70: S146. 10.1016/j.respe.2022.06.048

53. Piovezan R, Rosa SL, Rocha ML, de Azevedo TS, Von Zuben CJ. Entomological surveillance, spatial distribution, and diversity of Culicidae (Diptera) immatures in a rural area of the Atlantic Forest biome, State of São Paulo, Brazil. J Vector Ecol. 2013 Dec;38(2):317–25. doi: 10.1111/j.1948-7134.2013.12046.x. PMID: 24581361.

54. Juliano SA, O’Meara GF, Morrill JR, Cutwa MM. Desiccation and thermal tolerance of eggs and the coexistence of competing mosquitoes. Oecologia. 2002 Feb 1;130(3):458–469. doi: 10.1007/s004420100811. PMID: 20871747; PMCID: PMC2944657.

55. Weaver SC, Barrett AD. Transmission cycles, host range, evolution and emergence of arboviral disease. Nat Rev Microbiol. 2004 Oct;2(10):789–801. doi: 10.1038/nrmicro1006. PMID: 15378043; PMCID: PMC7097645.

56. Campos RK, Rossi SL, Tesh RB, Weaver SC. Zoonotic mosquito-borne arboviruses: Spillover, spillback, and realistic mitigation strategies. Sci Transl Med. 2023 Oct 18;15(718):eadj2166. doi: 10.1126/scitranslmed.adj2166. Epub 2023 Oct 18. PMID: 37851824; PMCID: PMC10807030.

57. Hanley KA, Cecilia H, Azar SR, Moehn BA, Gass JT, Oliveira da Silva NI, Yu W, Yun R, Althouse BM, Vasilakis N, Rossi SL. Trade-offs shaping transmission of sylvatic dengue and Zika viruses in monkey hosts. Nat Commun. 2024 Mar 27;15(1):2682. doi: 10.1038/s41467-024-46810-x. PMID: 38538621; PMCID: PMC10973334.

58. Diallo D, Diagne CT, Hanley KA, Sall AA, Buenemann M, Ba Y, Dia I, Weaver SC, Diallo M. Larval ecology of mosquitoes in sylvatic arbovirus foci in southeastern Senegal. Parasit Vectors. 2012 Dec 7;5:286. doi: 10.1186/1756-3305-5-286. PMID: 23216815; PMCID: PMC3543325.

59. Santana-Martínez JC, Molina J, Dussán J. Asymmetrical Competition between Aedes aegypti and Culex quinquefasciatus (Diptera: Culicidae) Coexisting in Breeding Sites. Insects. 2017 Oct 24;8(4):111. doi: 10.3390/insects8040111. PMID: 29064390; PMCID: PMC5746794.

60. Djamouko-Djonkam L, Nkahe DL, Kopya E, Talipouo A, Ngadjeu CS, Doumbe-Belisse P, Bamou R, Awono-Ambene P, Tchuinkam T, Wondji CS, Antonio-Nkondjio C. Implication of Anopheles funestus in malaria transmission in the city of Yaoundé, Cameroon. Parasite. 2020;27:10. doi: 10.1051/parasite/2020005. Epub 2020 Feb 12. PMID: 32048986; PMCID: PMC7015064.

61. Mathania MM, Munisi DZ, Silayo RS. Spatial and temporal distribution of *Anopheles* mosquito’s larvae and its determinants in two urban sites in Tanzania with different malaria transmission levels. Parasite Epidemiol Control. 2020 Sep 1;11:e00179. doi: 10.1016/j.parepi.2020.e00179. PMID: 32964148; PMCID: PMC7490549.

62. World Health Organization (WHO). WHO initiative to stop the spread of Anopheles stephensi in Africa, 2023 update WHO/UCN/GMP/2023.06. https://iris.who.int/bitstream/handle/10665/372259/WHO-UCN-GMP-2023.06-eng.pdf?sequence=1. Assessed 23 November 2024.

63. Taylor R, Messenger LA, Abeku TA, Clarke SE, Yadav RS, Lines J. Invasive Anopheles stephensi in Africa: insights from Asia. Trends Parasitol. 2024 Aug;40(8):731–743. doi: 10.1016/j.pt.2024.06.008. Epub 2024 Jul 24. PMID: 39054167.

64. Kamau WW, Sang R, Ogola EO, Rotich G, Getugi C, Agha SB, Menza N, Torto B, Tchouassi DP. Survival rate, blood feeding habits and sibling species composition of Aedes simpsoni complex: Implications for arbovirus transmission risk in East Africa. PLoS Negl Trop Dis. 2022 Jan 24;16(1):e0010171. doi: 10.1371/journal.pntd.0010171. PMID: 35073317; PMCID: PMC8812930.

65. Kamau WW, Sang R, Rotich G, Agha SB, Menza N, Torto B, et al. Patterns of Aedes aegypti abundance, survival, human-blood feeding and relationship with dengue risk, Kenya. Front Trop Dis. 2023;4: 1113531. doi:10.3389/fitd.2023.1113531.

66. Badolo A, Sombié A, Yaméogo F, Wangrawa DW, Sanon A, Pignatelli PM, Sanon A, Viana M, Kanuka H, Weetman D, McCall PJ. First comprehensive analysis of Aedes aegypti bionomics during an arbovirus outbreak in west Africa: Dengue in Ouagadougou, Burkina Faso, 2016-2017. PLoS Negl Trop Dis. 2022 Jul 6;16(7):e0010059. doi: 10.1371/journal.pntd.0010059. PMID: 35793379; PMCID: PMC9321428.

67. Agha SB, Tchouassi DP, Bastos ADS, Sang R. Assessment of risk of dengue and yellow fever virus transmission in three major Kenyan cities based on Stegomyia indices. PLoS Negl Trop Dis. 2017 Aug 17;11(8):e0005858. doi: 10.1371/journal.pntd.0005858. PMID: 28817563; PMCID: PMC5574621.

68. L’Azou M, Succo T, Kamagaté M, Ouattara A, Gilbernair E, Adjogoua E, Luxemburger C. Dengue: etiology of acute febrile illness in Abidjan, Côte d’Ivoire, in 2011-2012. Trans R Soc Trop Med Hyg. 2015 Nov;109(11):717–22. doi: 10.1093/trstmh/trv076. Epub 2015 Sep 18. PMID: 26385938; PMCID: PMC4603269.

69. Stoler J, Al Dashti R, Anto F, Fobil JN, Awandare GA. Deconstructing “malaria”: West Africa as the next front for dengue fever surveillance and control. Acta Trop. 2014 Jun;134:58–65. doi: 10.1016/j.actatropica.2014.02.017. Epub 2014 Mar 5. PMID: 24613157.

70. Saré D, Pérez D, Somé PA, Kafando Y, Barro A, Ridde V. Community-based dengue control intervention in Ouagadougou: intervention theory and implementation fidelity. Glob Health Res Policy. 2018 Aug 3;3:21. doi: 10.1186/s41256-018-0078-7. PMID: 30123837; PMCID: PMC6091010.

71. Bonnet E, Fournet F, Benmarhnia T, Ouedraogo S, Dabiré R, Ridde V. Impact of a community-based intervention on Aedes aegypti and its spatial distribution in Ouagadougou, Burkina Faso. Infect Dis Poverty. 2020 Jun 5;9(1):61. doi: 10.1186/s40249-020-00675-6. PMID: 32503665; PMCID: PMC7275586.

72. Sombié I, Degroote S, Somé PA, Ridde V. Analysis of the implementation of a community-based intervention to control dengue fever in Burkina Faso. Implement Sci. 2020 May 14;15(1):32. doi: 10.1186/s13012-020-00989-x. PMID: 32408903; PMCID: PMC7222308.

73. Zahouli JZB, Aboa PEB, Adjobi CN., Koffi V, Angoua ELE, Kouassa MA, Brika CA, Koua GK, Gbané A, Fofana D, Patel J, Thomas S, Leuenberger A, Vavassori L, Ruel-Bergeron S, Ferrari G, Müller P. Dengue vector control through multisectoral and community-based interventions in Abidjan, Côte d’Ivoire: study protocol for a cluster-randomised trial. Submitted to Trials.

